# Sprouty4 is required for Mdm2 regulation of invasion, focal adhesion formation and metastasis in cells lacking p53

**DOI:** 10.1101/2023.05.08.539890

**Authors:** Rafaela Muniz de Queiroz, Gizem Efe, Asja Guzman, Naoko Hashimoto, Yusuke Kawashima, Tomoaki Tanaka, Anil K Rustgi, Carol Prives

**Affiliations:** Department of Biological Sciences, Columbia University, New York, NY, USA 10027; Herbert Irving Comprehensive Cancer Center, Columbia University, New York, NY, USA 10032; Department of Molecular Diagnosis, Graduate School of Medicine, Chiba University, Chiba, Japan 260-8670; Department of Applied Genomics, Kazusa DNA Research Institute, Kisarazu, Chiba, Japan 292-0818; Research Institute of Disaster Medicine, Chiba University, Chiba, Japan

**Keywords:** Mdm2, p53-independent, focal adhesion, Sprouty4, migration, invasion, metastasis, cancer

## Abstract

Although the E3 ligase Mdm2 and its homologue and binding partner MdmX are the major regulators of the p53 tumor suppressor protein, it is now evident that Mdm2 and MdmX have multiple functions that are independent of p53. For example, Mdm2 can regulate cell migration, although mechanistic insight into this function is still lacking. Here we show in cells lacking p53 expression that knockdown of Mdm2 or MdmX, as well as pharmacological inhibition of the Mdm2/MdmX complex, not only reduces cell migration and invasion, but also impairs cell spreading and focal adhesion formation. In addition, Mdm2 knockdown decreases metastasis *in vivo*. Remarkably, Mdm2 modulates the expression of Sprouty4, which is required for the Mdm2 mediated effects on cell migration, focal adhesion formation and metastasis. Our findings describe a molecular mechanism by which the Mdm2-X complex, through Sprouty4, regulates cellular processes leading to decreased metastatic capability independent of p53.

## Introduction

Mouse Double Minute 2 (Mdm2) was first described as a potential oncogene in mouse fibroblasts ^1^. Shortly thereafter Mdm2 was discovered to bind the p53 tumor suppressor protein, which was followed by studies validating it as the most critical and obligate regulator of p53 ^2–5^. Mdm2 is an E3 ligase that can function either alone or with its homologue MdmX (aka Mdm4) in a complex to ubiquitinate, SUMOylate or NEDDylate proteins, most notably p53^3, 6^.

While *TP53* is found mutated in more than 50% of all human cancers ^7^, *Mdm2* is rarely lost or deleted in cancer. Instead, the *Mdm2* gene is found amplified in a number of tumors, notably sarcomas, but also in melanomas, glioblastomas and breast cancers ^8^. *MdmX* is also over-expressed in several cancers, both where *Mdm2* is also amplified as well as others, such as retinoblastomas and hematological cancers, where *MdmX* alone is amplified^9, 10^.

Although the great majority of published studies has focused on the relationship between Mdm2, MdmX and p53, there is a newly literature describing p53-independent activities of Mdm2 and MdmX^6, 11^. Indeed, Mdm2 has been shown to regulate many cellular processes involved in tumorigenesis such as regulation of chromatin state^12^ and DNA repair^11^, maintenance of cell survival and growth^13, 14^, modulation of cell migration^15, 16^ and promotion of metastasis^17, 18^. In aggregate the abovementioned findings have reintroduced Mdm2 as a potential oncogene in its own right, as first suggested by Fakharzadeh in 1991

Mdm2 regulation of cell migration has revealed that post-translational modifications and microRNAs play roles ^16, 19–21^, but underlying mechanisms remain to be elucidated. Most reported experiments testing migration/invasion were performed in “2D” settings (i.e. cells attached to and growing on plastic culture dishes) as opposed to the “3D” model where cells are placed on or within a substratum that resembles the extracellular matrix (ECM) and form clusters that more closely recapitulate the *bona fide* tissue architecture. Further, previous studies describing changes in cell migration have only explored the impact of altering Mdm2 protein levels and did not address whether the E3-ligase activity of Mdm2 and the Mdm2/MdmX complex are required for this function. The same can be said for studies examining the potential roles of Mdm2 in metastasis ^17, 18, 22, 23^.

One additional confounding issue of previous studies characterizing pro-oncogenic activities of Mdm2 has been that for the most part researchers have used cell lines that either harbor wild-type or mutant versions of p53 to make their observations. While in such cases modulation of Mdm2 levels in different cell lines harboring either wild-type or different mutant versions of p53 leads to changes in cell migration^15, 20, 21, 24–27^, the presence of either form of p53 might impact cellular responses, however, and thereby complicate interpretation of data. Here, by both altering Mdm2 levels and inhibiting the E3 ligase activity of the Mdm2/MdmX complex in cells that lack p53 protein, we have been able to delve into fully p53-independent mechanism(s) by which Mdm2 regulates migration and invasion *in vitro* and metastasis *in vivo*.

## Results

### Mdm2 regulates cell migration, cell attachment to extracellular matrix components and cell spreading independently of p53 and EMT

To eliminate possible complications in data interpretation stemming from the presence of p53 protein, we used both a CRISPR-derived human HT1080 fibrosarcoma p53 knock-out cell line (HT1080 p53KO; Supplementary Fig. 1A) that does not express p53 and the H1299 lung carcinoma-derived cell line that is endogenously p53-null. While it was previously reported that lowering Mdm2 levels typically with either si- or sh-RNAs, interferes with cell migration^23, 28, 29^, to our knowledge the importance of the E3-ligase activity of Mdm2 and the formation of the Mdm2-X complex in this process has not been reported. To test this, together with Mdm2 or MdmX silencing using siRNAs, we have used a small molecule inhibitor of Mdm2, MEL23, that was previously shown to inhibit the formation and function of the Mdm2/MdmX complex, leading to accumulation of Mdm2 and MdmX proteins ^30^.

We treated HT1080 p53KO cells either with Mdm2 siRNAs (Fig. 1A), or with MEL23, which, as expected, led to increased Mdm2 and MdmX protein levels (Fig. 1B). Cells were also treated with MdmX siRNAs (Supplementary Fig. 1B). Either siRNA-mediated knockdown of Mdm2 (Fig. 1C), MEL 23 treatment (Fig. 1D) or MdmX siRNAs (Supplementary Fig. 1C) led to reduced cell migration using the wound scratch assay.

**Fig. 1.**
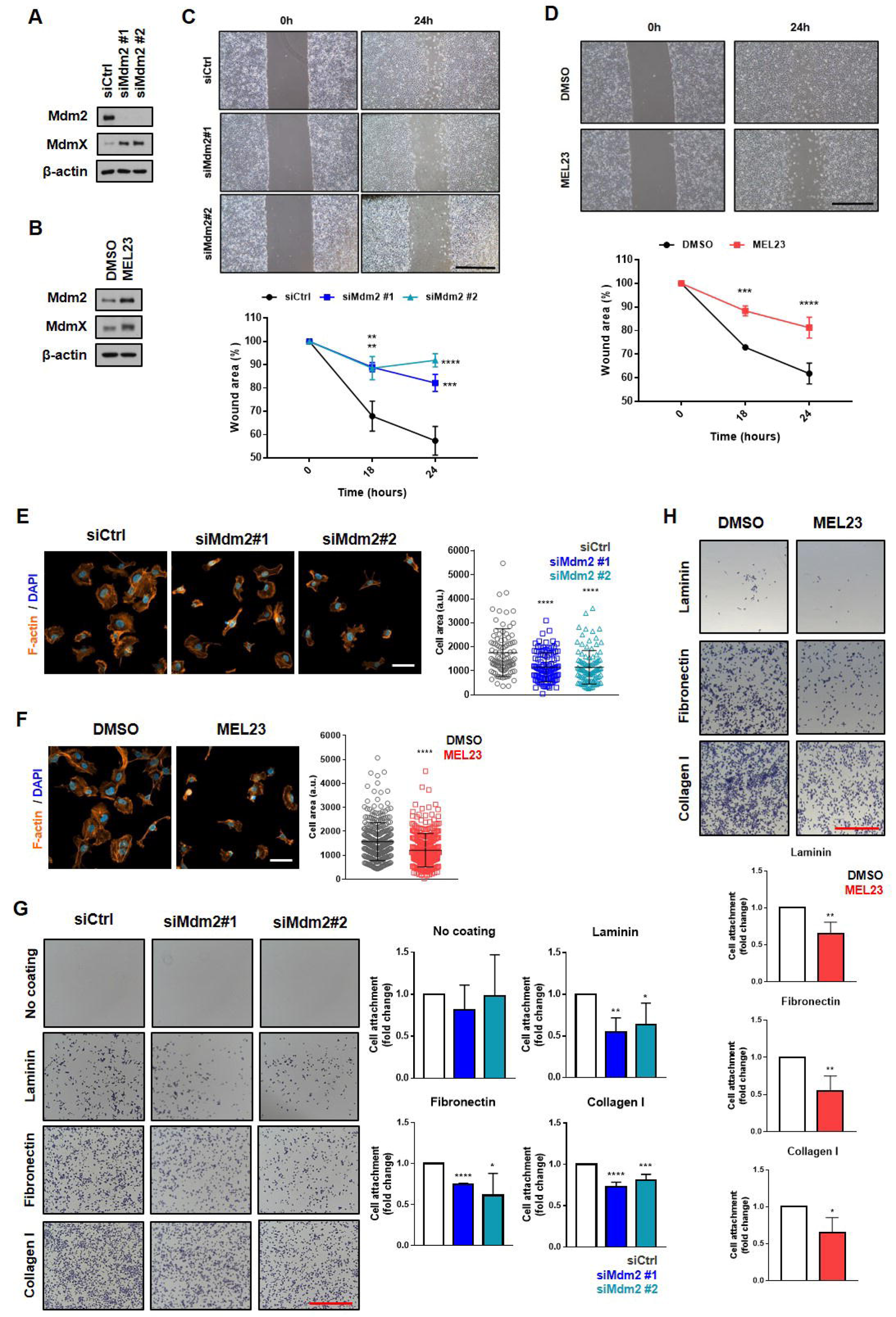
Loss of Mdm2 or functional impairment of the Mdm2/MdmX complex decreases migration, impairs cell spreading and cell attachment to ECM in a p53-independent manner. HT1080 p53KO cells were transfected with siRNAs against Mdm2 and siCtrl or treated with the Mdm2 inhibitor, MEL23. (A-B) Protein levels of Mdm2 and MdmX after (A) transfection using siRNAs as indicated or (B) treatment with MEL23 (7 µM) for 24 h. β-actin was used as loading control. (C-D) Cell migration assay. Representative images (top) and quantification (bottom) of wound scratch migration assay with cells (C) silenced for Mdm2 or (D) treated with MEL23 for 24 h. Scale bar equivalent to 1 mm. (E-F) Representative images of morphology of cells attached to collagen coated coverslips and quantification of cell area after (E) silencing of Mdm2 or (F) treatment with MEL23 for 24 h. Cells stained for actin (orange) and DNA via DAPI (blue). Scale bar equivalent to 50 µm. (G-H) Quantification and representative micrographs showing attachment to different ECM components of cells (G) silenced for Mdm2 or (H) treated with MEL23 for 24 h. Scale bar equivalent to 1 mm. Experiments shown represent mean ±SD of at least 3 biological replicates, *p<0.05, **p<0.01, ***p<0.001, ****p<0.0001.

When Mdm2 or MdmX was silenced by siRNA or when the complex was inhibited by MEL23, the morphology of cells migrating into the wound during migration assays changed. Thus, we analyzed the morphology of cells in 2D in response to those stimuli and we found that the area of attaching cells was significantly reduced (Fig. 1E and F and Supplementary Fig 1D), while in solution (unattached) cell size did not change after such treatments (Supplementary Fig. 1E) demonstrating the changes in area are not due to cell volume.

Since the ability of cells to spread is correlated directly to interact and bind to molecules in the extracellular environment, we performed an attachment assay using the three main extracellular matrix (ECM) components, collagen (represented by collagen I), fibronectin, and laminin. When Mdm2 was silenced by siRNA or functionally inhibited by MEL23, the attachment capability of the HT1080 p53KO cells was significantly reduced to a very similar extent for all three ECM components (Fig. 1G and H). It should be pointed out that although Mdm2 modulation has been associated with Epithelial-Mesenchymal Transition/Mesenchymal-Epithelial Transition (EMT/MET) in some cells^17, 23, 31, 32^, there were no changes associated with different EMT markers (E-cadherin, N-cadherin, vimentin and Zeb1) in response to silencing Mdm2 (Supplementary Fig. 1F).

Taken together, our results support the likelihood that the active Mdm2/MdmX complex is required for efficient migration of cells growing in 2D culture conditions, which likely results from impaired ability to interact with or respond to the ECM.

### Functional impairment of the Mdm2/X complex decreases individual and collective invasion in a 3D *in vitro* model of cancer cell dissemination

Experimental results related to cell migration often differ substantially between 2D and 3D models ^33, 34^. To gain insights on the impact of manipulation of Mdm2 levels or activity in a 3D setting, we examined individual and collective invasion of HT1080 KO cells using a physiologically relevant 3D tumor spheroid model ^35^. Spheroid invasion in a low density collagen I matrix was used to assess individual cancer cell invasion, since it is a more permissive matrix, while invasion in a more dense composite matrix (collagen I/BME, Basement Membrane Extract) was used to assess collective invasion ^36^. We found that silencing of Mdm2 as well as treatment with MEL23 led to a significant decrease in individual 3D invasion, which is typically determined by measuring the area invaded over a given time period and by counting the number of cells invading from the spheroid into the surrounding ECM (Fig. 2A and B). When using the biomechanically more challenging composite matrix (collagen/BME), we observed that both siRNA and MEL23 treatment significantly decreased collective invasion as well (Fig. 2C and D). We also embedded dispersed cells in collagen I matrices to investigate if cell morphology in this 3D setting was affected by Mdm2 knockdown, as we had observed in 2D cultures. Again, silencing of Mdm2 led to altered cell morphology; specifically, there was a statistically significant shift towards a rounder cell morphology (cells with higher circularity) in 3D, when compared to the control group (Fig. 2E and F). It should be noted that Mdm2 knockdown did not affect spheroid proliferation for the duration of the invasion assay (Supplementary Fig. 2A). Our results support a requirement for functional Mdm2 to serve as a key regulator of individual and collective cell invasion. They provide a premise in which to examine whether Mdm2 might play a role in processes associated with metastasis *in vivo*.

**Fig. 2.**
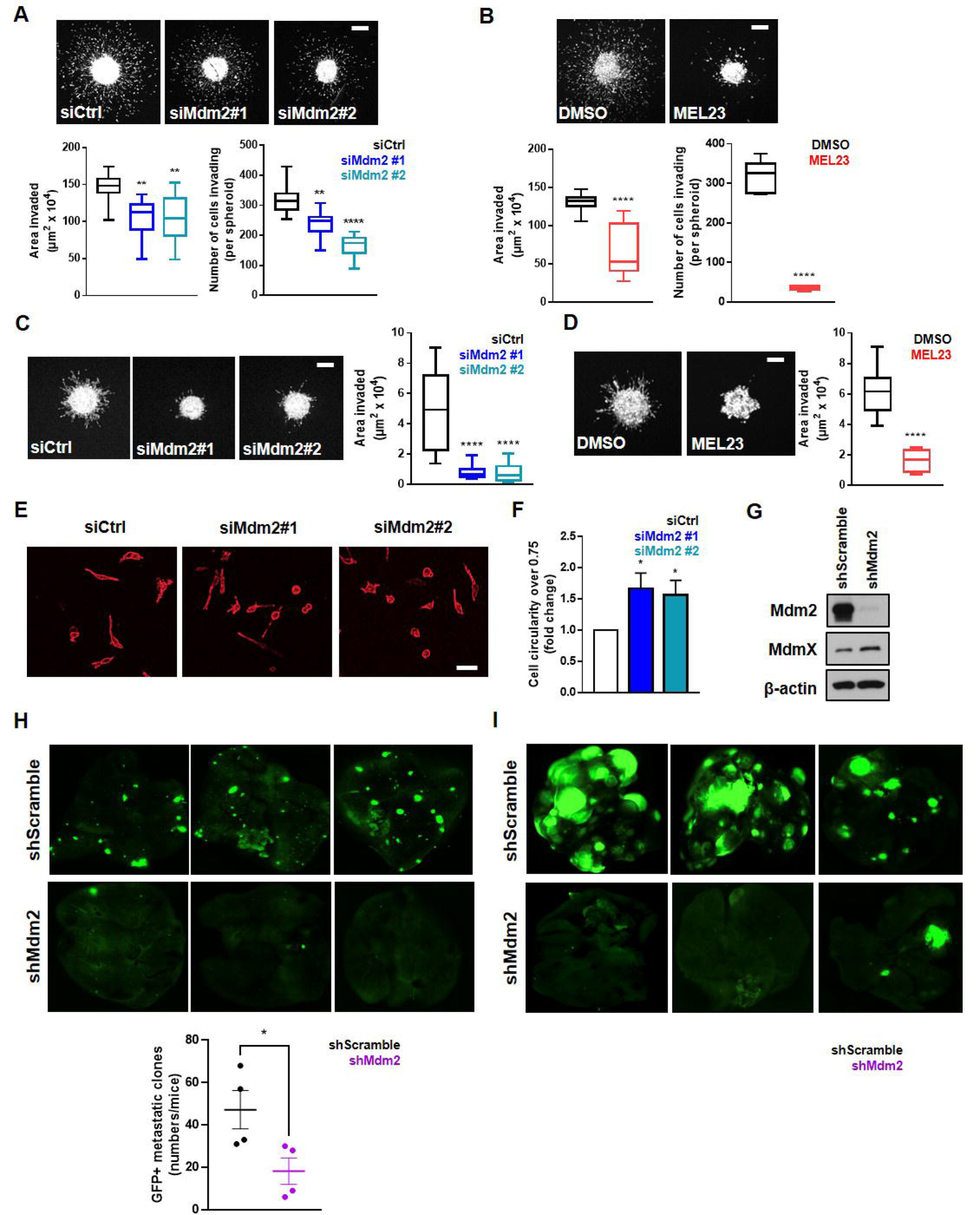
Loss of Mdm2 function decreases individual and collective invasion in 3D and metastatic burden *in vivo*. **(A-B)** Tumor spheroid invasion in collagen I matrix after (A) transfection of HT1080 p53KO cells with siRNAs against Mdm2 or (B) treatment with MEL23. Representative images above and quantification below of area invaded and number of cells invading 24 h after implantation. **(C-D)** Tumor spheroid invasion in collagen I-BME matrix after (C) transfection of HT1080 p53KO cells with siRNAs against Mdm2 or (D) treatment with MEL23. Representative images and quantification of area invaded 24 h after implantation. **(E)** Representative images of cell morphology of HT1080 p53KO cells transfected with siRNAs against Mdm2 or siCtrl in collagen matrix, 6 h after implantation. **(F)** Quantification of more circular cells (circularity of 0.75 or higher) in each condition shown in (E). **(G-I)** HT1080 p53KO cells stably expressing shRNA scramble (shScramble) or a pool of shRNAs against Mdm2 (shMdm2) were used to analyze metastatic burden using mouse models. (G) Protein levels of Mdm2 and MdmX in HT1080 shScramble and shMdm2 stable cell lines. β-actin was used as loading control. (H) Representative images above and quantification below of metastatic foci in the lungs after implantation of shScramble or shMdm2 cells using orthotropic model. (I) Representative images above and quantification below of metastatic foci in the lungs after injection of shScramble or shMdm2 cells using tail-vein model. Scale bars equivalent to 200 µm. Experiments shown represent mean ±SD of at least 3 biological replicates, *p<0.05, **p<0.01, ***p<0.001, ****p<0.0001.

In summary, when one or the other subunit of the Mdm2/X complex is depleted, as well as in the presence of both proteins that cannot form an active E3-ligase, the cells we studied display impaired migration and attachment as well as invasion. These findings indicate that a functional Mdm2/X complex is needed for the regulation of cell migration and invasion by Mdm2.

### Mdm2 knockdown decreases metastatic burden *in vivo*

To evaluate if decreased Mdm2 levels can impact metastasis we performed experiments using both orthotopic and tail-vein injections in athymic nude mice. We first established GFP-expressing cell lines derived from HT1080 p53KO cells that stably express either normal (shScramble cells) or much lower Mdm2 levels (shMdm2 cells that harbored four different shRNAs) (Fig. 2G). Assessment of cell motility (Supplementary Fig. 2B) and cell morphology (Supplementary Fig. 2C) demonstrated that the HT1080 cell line with stable Mdm2 knockdown (shMdm2) displayed the same characteristics observed in cells transiently transfected with siRNA or in cells treated with MEL23 when compared to the shScramble cells.

Using these GFP^+^ HT1080 p53KO cell lines, we performed an analysis of metastatic burden in the lungs first using an orthotopic model *in vivo* where cells were introduced into the quadriceps femoris muscle of mice, and cells from tumors formed *in situ* were allowed to metastasize to the lungs. After 6-weeks (counting from the injection of tumor cells) mice were euthanized and the number of metastatic foci in the lungs were analyzed. The mice injected with shMdm2 cells Mdm2 had a dramatic and significant decrease in the number of metastatic foci in comparison to the control shScramble group (Fig. 2H). Measurement of tumor weight in our orthotopic model showed that tumors with Mdm2 knocked down were significantly smaller than in the control group at the time of harvest (Supplementary Fig. 2D). While these findings revealed that reducing Mdm2 levels impacted metastasis, we could not rule out the possibility that at least some of the effects seen were due to the initial smaller sized shMdm2 tumors that formed over the course of these experiments.

To better understand if the changes in metastatic burden observed in mice were due to decreased metastatic ability or changes in primary tumor growth between the two groups, we performed lateral tail-vein injections into athymic nude mice. This model encompasses the stages of metastatic cascade after the tumor cells enter the bloodstream, and is not influenced by differences in initial tumor size, since the same number of cells for the two experimental groups are injected into the mice. The analysis of metastatic burden in the lungs of mice injected with the GFP^+^ HT1080 p53 KO cells using this model demonstrated once again that cells expressing less Mdm2 had reduced metastatic capability in comparison to the control group *in vivo* (Fig. 2I). These data suggest that defects in cell attachment and motility of cancer cells lacking full levels of Mdm2 may lead to a decreased ability of cancer cells to metastasize into distant organs.

### Mdm2 depletion or inhibition decreases number and size of focal adhesions

Our findings that knockdown or pharmacological inhibition of Mdm2 decreased cell spreading and cell attachment to the ECM accompanied by reduced migration, invasion and metastasis, suggested that this might be due to inhibition of one or more signaling processes responsible for intermediating cellular contacts with the extracellular environment. It is well established that the main proteins that mediate ECM attachment are integrins, that each have specific sets of ECM substrates^37^. Yet, since those ECM substrates that we tested were affected to similar extents, we speculated that integrins are not the likely step in the process of ECM attachment that is affected by Mdm2 downregulation. In concert with this, when we measured the levels of those integrins by mass spectrometry (which is described in the next section) or the most expressed integrins in HT1080 cells (integrin beta 1, integrin alpha 2 and integrin alpha 3) by immunoblot we did not identify any that were significantly affected by Mdm2 knockdown or inhibition (Supplementary Table 1 and Supplementary Fig 3A).

Activation of integrins leads to recruitment of a set of proteins that form structures known as focal adhesions (FA), which are responsible for integrin activation via linking cells to the ECM with ensuing reorganization of the cytoskeleton in order to promote anchorage and support motility ^38, 39^. Strikingly, when HT1080 cells were transfected with Mdm2 siRNAs (Fig. 3A), treated with MEL23 (Fig. 3B) or transfected with MdmX siRNAs (Supplementary Fig. 3B), the number and size of FA foci decreased significantly. Notably, the stable cell line expressing shMdm2 used in the metastasis experiments described above also displayed reduced FAs compared to control cells (Supplementary Fig. 3C).

**Fig. 3.**
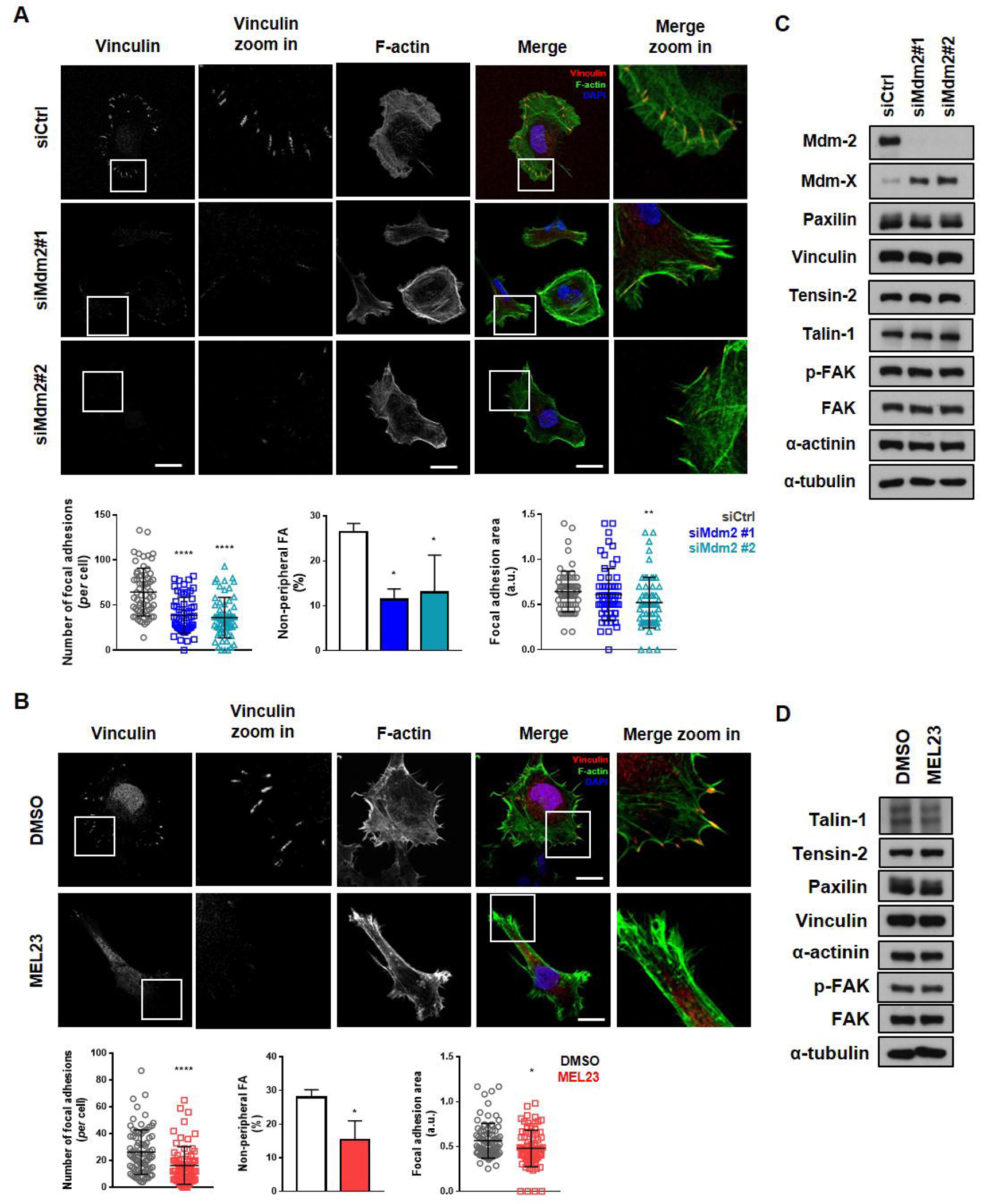
Loss of Mdm2 or functional impairment of Mdm2/MdmX complex decreases FA formation in a p53-independent manner. HT1080 p53KO cells were (A) silenced for Mdm2 using siRNAs or (B) treated with 7 µM MEL23 for 24 h. **(A-B)** Immunofluorescence showing FA foci by vinculin staining (red), stress fiber formation by phalloidin staining of F-actin (green), and nuclei (blue) detected by DAPI staining of DNA. Representative images shown above with quantification of FA parameters below. Scale bar equivalent to 20 µm. **(C-D)** Protein levels of FA related proteins in HT1080 p53KO cells (C) silenced for Mdm2 or (D) treated with MEL23 for 24 h. α-tubulin was used as loading control. Experiments shown represent mean ±SD of at least 3 biological replicates, *p<0.05, **p<0.01, ***p<0.001, ****p<0.0001.

The process of FA formation starts with transient structures called nascent adhesions (NAs). These are formed by integrin activation and recruitment of FA-related proteins, such as talin, paxilin and vinculin. These NAs then cluster in order to mature into FAs that then move inwards and can further mature to form focal complexes (FC) (also known as fibrilar adhesions). If, upon integrin activation, NAs don’t cluster, the structure is then disassembled ^40^. The process of FA maturation is not yet fully understood but the size and the subcellular localization of a NAs, FAs or FCs, as well as their linkage to stress fibers, can be used to indicate the maturation state of the structure ^40, 41^. Upon Mdm2 silencing FA foci were not only reduced in number, but their relative size was reduced (Fig. 3A and B right graphs) along with fewer non-peripheral FAs (Fig. 3A and B center graphs). Moreover, F-actin staining to probe stress fibers showed a much denser F-actin structure in control conditions compared to cells treated with Mdm2 siRNA and MEL23 (Fig. 3A and B, F-actin column). Together, our results indicate that Mdm2 plays a role in the maturation of NAs into FAs and FCs.

### The Mdm2-MdmX complex regulates Sprouty4 expression

Although we had linked Mdm2 functionally with FA formation, when we probed the levels of several proteins known to be involved in the formation of FAs, somewhat confoundingly none of those proteins tested were significantly altered in response to Mdm2 knockdown or MEL23 treatment (Fig. 3C and D). As a result, we went on to perform an untargeted proteomic analysis of cells treated with MEL23 or transfected with siRNA against Mdm2. Validation of the conditions used was reflected by levels of Mdm2 that were in accordance with both the immunoblot analysis of proteomics’ samples and the results of the proteomics analysis itself (Supplementary Fig 4A and B). As expected, many proteins in both groups (siRNA and MEL23) were significantly up- or downregulated after these treatments. The Volcano plots show all identified peptides and their significance based on statistical analysis (Fig. 4A).

**Fig. 4.**
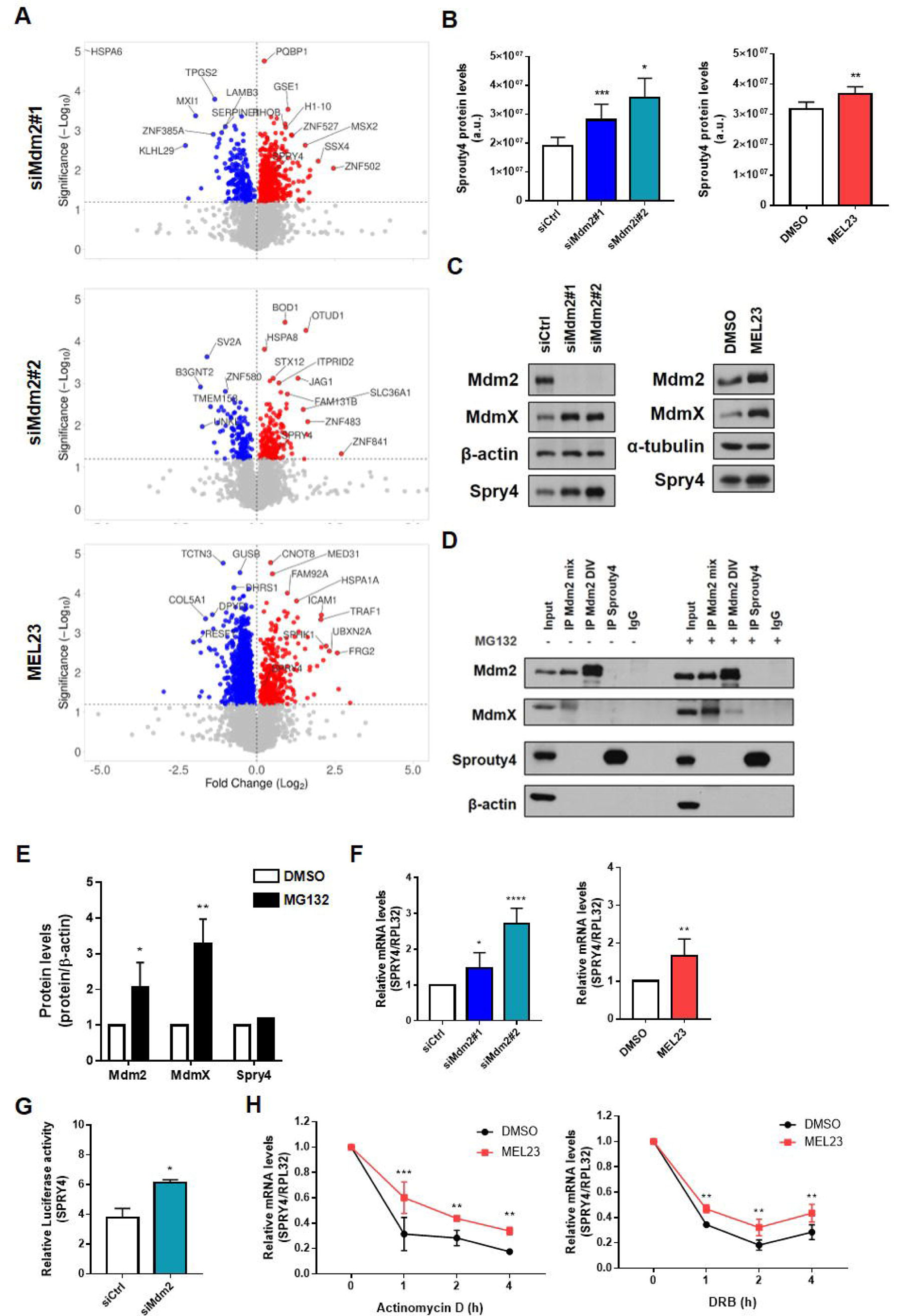
The Mdm2/MdmX complex regulates expression of Sprouty4. HT1080 p53KO cells were transfected with siRNAs against Mdm2 or siCtrl for 24 h; or treated with 7 µM MEL23 or DMSO (vehicle) for 24 h. **(A)** Volcano plots show all peptides identified by mass spectrometry. Colored dots represent significantly differentially expressed proteins that were downregulated (blue dots) and upregulated (red dots) in each condition shown at left of plot. Gray dots represent non-significant changes. MEL23 treated cells were compared to DMSO treated cells. Cells transfected with siMdm2#1 or #2 were compared to siCtrl-transfected cells. **(B-C)** Spry4 expression in HT1080 p53KO cells in response to Mdm2 knockdown or treatment with MEL23 for 24 h by (B) mass spectrometry or (C) immunoblotting. β-actin and α-tubulin were used as loading control for immunoblot. **(D)** Co-immunoprecipitation of Mdm2 and Spry4 in the presence or absence of MG132. β-actin was used as loading control. **(E)** Quantification of Spry4, Mdm2 and MdmX protein levels after treatment with MG132 (or vehicle, DMSO) for 4 h. **(F)** Spry4 mRNA levels in response to Mdm2 knockdown or treatment with MEL23 for 24 h. RPL32 was used as housekeeping. **(G)** *Spry4* luciferase promoter assay in cells silenced for Mdm2 using siRNA. **(H)** Spry4 mRNA levels in cells treated with MEL23 or vehicle at the indicated times after treatment with DMSO, ActinomycinD (10 µg/ml), or DRB (100 µM) as indicated. Experiments shown represent mean ±SD of at least 3 biological replicates, *p<0.05, **p<0.01, ***p<0.001.

When we compared significantly differentially expressed proteins between the different conditions in the two groups showed that in the siRNA group (siRNA#1 and siRNA#2 vs. siCtrl), we found 799 differentially expressed proteins, while in the drug group (MEL23 vs. DMSO) 1854 proteins were differentially expressed (see heat maps in Supplementary Fig. 4C). From these two datasets 146 proteins were significantly differentially expressed by both siRNA and MEL23 (Supplementary Fig 4D). Pathway analysis of those 146 proteins revealed that among those most impacted by siMdm2 or MEL23 were signaling pathways involved in cell adhesion, namely those involved in focal adhesion (MAPK, Ras, PI3K and GPCR) (Supplementary Fig. 4E).

After assessing those differentially expressed 146 proteins and focusing on those associated with cell migration and motility, Sprouty4 (Spry4) emerged as a viable candidate for further investigation. Spry4 is one of four members of the Sprouty family (which includes Spry 1, 2, 3 and 4 as well as Spred1 and 2) that have been shown to be key negative regulators of Ras/ERK signaling. The expression and cancer relevance of different Sprouty proteins varies amongst different tumor types ^42, 43^. This was supported by our analysis of TCGA and GTEX databases where expression of the different Sprouty family members varies in different tissues, as well as when tumors are compared to normal tissue (Supplementary Fig. 5A), suggesting that they play different roles in tumorigenesis depending on the context. The same is seen for their potential value as prognostic markers ^42, 43^.

In accordance with the differences seen in the proteomics analysis (Fig. 4B) Spry4 levels were significantly increased (∼1.5 to 2-fold) in HT1080 p53KO cells that were treated with either siMdm2 or MEL23 (Fig. 4C and Supplementary Fig. 5B) measured by immunoblot. Furthermore, HT1080 shMdm2 cells as well as H1299 lung adenocarcinoma cells displayed increased Spry4 levels upon Mdm2 downregulation (Supplementary Fig. 5C and D). Although HT1080 cells express other Sprouty family members including Spry2 as well as Spred1 and 2, these were not differentially expressed in response to Mdm2 siRNAs or MEL23 treatment as determined by the proteomics analysis (Supplementary Fig. 5E).

We continued to focus on Spry4 and asked whether Mdm2 or the Mdm2 complex directly regulates Spry4 protein stability. As expected, MdmX was co-immunoprecipitated with Mdm2, however Spry4 was not pulled down with Mdm2 and the reverse co-immunoprecipitation provided the same negative result (Fig. 4D).

Further, cells treated with the proteasome inhibitor MG132 revealed no change in Spry4 protein levels in comparison to non-treated cells (Fig. 4D), which we confirmed by densitometry in all biological replicates (Fig. 4E). By contrast, as expected, Mdm2 and MdmX, both known targets of the proteasome, accumulated in the presence of MG132 (Fig. 4D and E).

We then determined that there was a significant increase in Spry4 mRNA in cells harboring siMdm2 or treated with MEL23 (Fig. 4F). To understand which aspect of Spry4 mRNA expression was being regulated by Mdm2 we tested both a Spry4 promoter luciferase reporter and determined Spry4 mRNA stability. Surprisingly, both Spry4 transcription (as determined by the reporter assay) (Fig 4G) and mRNA stability, as measured using two different transcription inhibitors (Fig. 4H and Supplementary Fig. 5F), were increased upon Mdm2 inhibition.

Finally, to confirm that the results are not unique to HT1080 cells we showed that knockdown of Mdm2 in H1299 cells (Fig. 5A) led to: decreased cell migration (Fig. 5B), decreased attachment to collagen (Fig. 5C) and impaired FA formation (Fig. 5D). As seen in HT1080 cells, Mdm2 silencing induces Spry4 expression at the protein and mRNA levels (Fig. 5E and F) in H1299 cells, thereby extending our results to a cell line which had evolved naturally to lack expression of p53. Most relevantly to our work, analysis of patient samples from TCGA and GTEX databases revealed that Spry4 expression is decreased in lung cancers, both squamous and adenocarcinoma, compared to normal lung tissues (Supplementary Fig 5G).

**Fig. 5.**
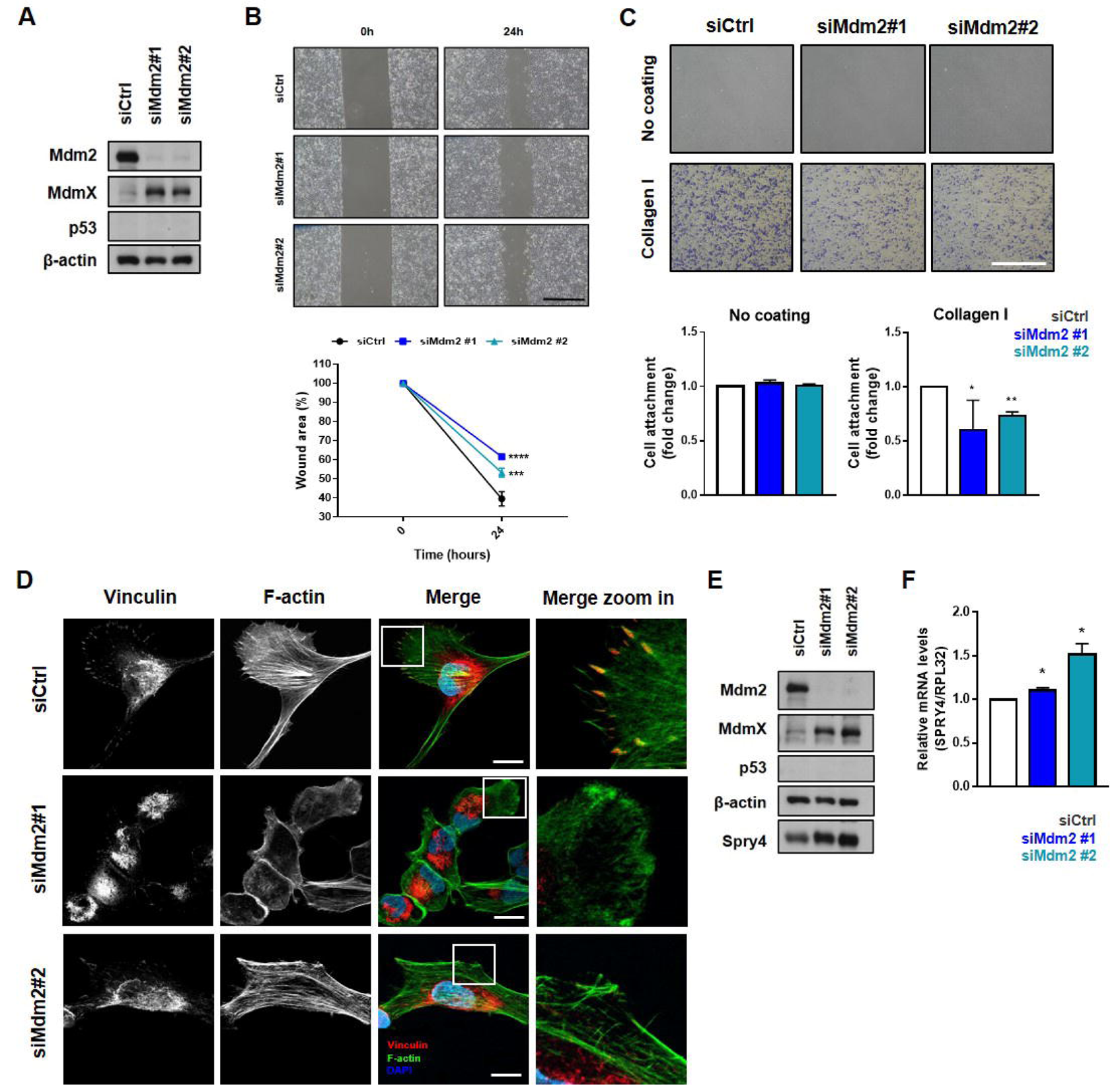
Mdm2 knockdown decreases cell migration, attachment to the ECM and FA formation while induces expression of Sprouty4 in naturally p53 null lung adenocarcinoma cell line. H1299 cells were transfected with siRNAs against Mdm2 and siCtrl**. (A)** Protein levels of Mdm2, MdmX and p53 after. β-actin was used as loading control. **(B)** Cell migration assay. Representative images (top) and quantification (bottom) of wound scratch migration assay. Scale bar equivalent to 1 mm. **(C)** Quantification and representative micrographs showing attachment to ECM component, collagen I. Scale bar equivalent to 1 mm. **(D)** Representative images of immunofluorescence showing FA foci by vinculin staining (red), stress fiber formation by phalloidin staining of F-actin (green), and nuclei (blue) detected by DAPI staining of DNA. Scale bar equivalent to 20 µm. **(E)** Protein levels of Spry4 as well as Mdm2, MdmX and p53 after Mdm2 silencing using siRNAs. β-actin was used as loading control. **(F)** mRNA levels of Spry4 after Mdm2 silencing using siRNAs. RPL32 was used as housekeeping. Experiments shown represent mean ±SD of at least 3 biological replicates, *p<0.05, **p<0.01, ****p<0.0001.

### Sprouty4 is required for Mdm2 to promote FA formation and metastasis

Overexpression of Spry4 has been reported to decrease migration of tumor cells ^44–47^. In order to understand if the noted increase in Spry4 protein levels was linked to changes in cell migration induced by Mdm2 inhibition, we tested the impact of reducing Spry4 under these experimental conditions. Immunoblotting showed that in the double knockdown (siMdm2+siSpry4) both Mdm2 and Spry 4 levels were greatly diminished (Fig. 6A), although cell viability was unaffected by depletion of either Mdm2 and Spry4 individually or together (Supplementary Fig. 6A). Analysis of cell migration under the same conditions revealed that co-silencing of Mdm2 and Spry4 completely rescued the reduction in cell migration triggered by Mdm2 silencing alone (Fig. 6B). Spry4 knockdown also rescued migration in MEL23-treated HT1080 p53KO cells (Supplementary Fig.6B). Co-depletion of Mdm2 and Spry4 also rescued the impaired cell spreading seen when Mdm2 alone was silenced, as shown by the restoration of HT1080 p53 KO cells’ area (Fig. 6C). Importantly, the double knockdown (siMdm2+siSpry4) led to the same FA pattern that was observed in the control condition (siCtrl), showing that ablation of Spry4 significantly rescued the FA impairment induced by siMdm2 (Fig. 6D).

**Fig. 6.**
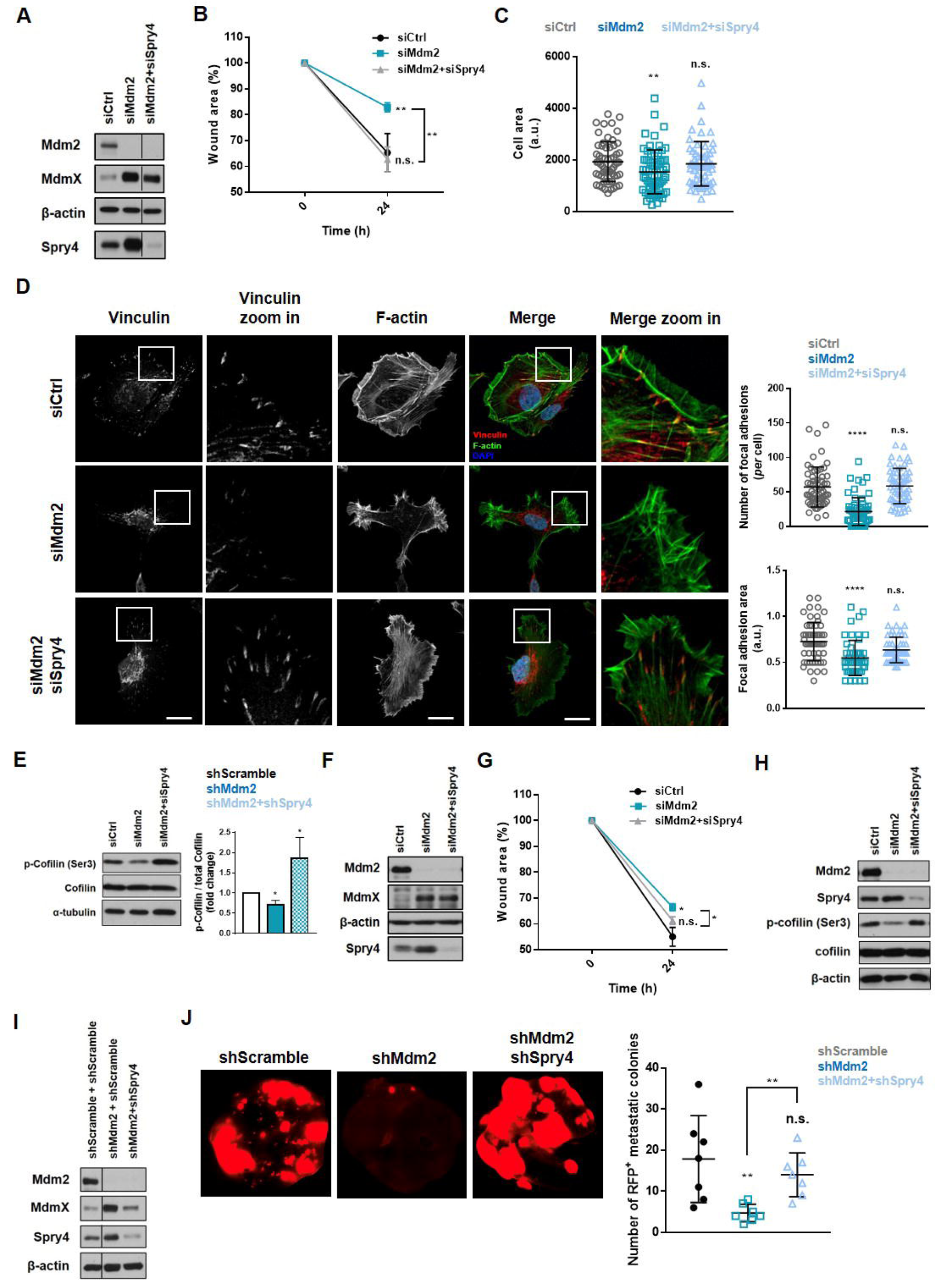
Knockdown of Sprouty4 rescues changes in cell migration, FA formation and metastasis resulting from loss of Mdm2. **(A-E)** HT1080 p53KO cells were transfected with siRNA against Mdm2 alone or both siRNA against Mdm2 and a pool of Spry4 siRNAs. (A) Protein levels of Mdm2, MdmX and Spry4 after transfection with indicated siRNAs. β-actin was used as loading control. (B) Quantification of wound scratch migration assay comparing migration into wound scratches in cells treated with the indicated siRNAs as in Fig. 1C. (C) Quantification of cell area after attachment to collagen coated coverslips as in Fig. 1E. (D) Immunofluorescence showing FA foci by vinculin staining (red), stress fiber formation by phalloidin staining of F-actin (green), and nuclei (blue) as detected by DAPI staining of DNA. Representative images shown at left and quantification of FA parameters shown at right. Scale bar equivalent to 20 µm. (E) Immunoblot (left) and densitometry (right) of levels of total and phospho-cofilin-1(Ser3) in HT1080 p53KO cells silenced for Mdm2 alone or with double KD of Mdm2 and Spry4. **(F-H)** H1299 cells were transfected with siRNA against Mdm2 alone or both siRNA against Mdm2 and a pool of Spry4 siRNAs. (F) Protein levels of Mdm2, MdmX and Spry4 after transfection with indicated siRNAs. β-actin was used as loading control. (G) Quantification of wound scratch migration assay comparing migration into wound scratches in cells treated with the indicated siRNAs as in Fig. 1C. (H) Immunoblot of levels of total and phospho-cofilin-1(Ser3) in H1299 cells silenced for Mdm2 alone or with double KD of Mdm2 and Spry4. **(I-J)** HT1080 p53KO cell lines were established stably expressing a pool of shRNAs against Mdm2 alone or Mdm2 and Spry4 together. (I) Protein levels of Mdm2, MdmX and Spry4 in stable cells lines. β-actin was used as loading control. (J) Analysis of metastatic burden *in vivo* using tail-vein injection model as in Fig.3E and 3F. Representative images above and quantification below of metastatic foci in the lungs after 8 weeks of injection. Experiments shown represent mean ±SD of at least 3 biological replicates, *p<0.05, **p<0.01, ****p<0.0001, n.s.: not significant.

As mentioned, Sprouty family members are most well characterized as negative regulators of Ras/ERK signaling^42^. To understand if this is the pathway affected by Mdm2 silencing we measured ERK phosphorylation in HT1080 p53KO cells. Surprisingly, when cells were silenced for Mdm2 there was no detectable impact on ERK phosphorylation, nor were there changes in ERK phosphorylation when both Mdm2 and Spry4 were depleted when compared to siMdm2 or siCtrl (Supplementary Fig. 6C). This indicates that Spry4 might act through a non-canonical pathway to induce the changes in cell migration in response to Mdm2 inhibition.

Spry4 has been reported previously to regulate phosphorylation of Cofilin-1 ^48^, an important event in actin filament dynamics that is involved in FA stability ^49^. Indeed, we observed that Mdm2 knockdown significantly decreased Cofilin-1 phosphorylation, and this was rescued by the co-knockdown of Mdm2 and Spry4 in HT1080 p53KO cells (Fig. 6 E). Staining for filamented (F-actin) and monomeric (G-actin) actins showed that silencing of Mdm2 favored monomeric actin, which was reversed in the double knockdown condition (Supplementary Fig. 6D), confirming our observations of altered cofilin phosphorylation. Double knockdown of Mdm2 and Spry4 also rescued migration in H1299 cells (Fig. 6F and G) as well as the phosphorylation of cofilin, which is also reduced after Mdm2 knockdown alone (Fig. 6H).

We then turned to determine whether the regulation of Spry4 expression by Mdm2 might be involved in the role of Mdm2 in metastasis *in vivo*. For this we generated cells stably co-expressing shRNAs against Mdm2 and Spry4 for comparing with the cells we had generated expressing shRNAs vs. Mdm2 alone. Confirmation of the silencing of Spry4 and Mdm2 in these stable cells lines was measured by immunoblotting (Fig. 6I) and a migration assay confirmed that the stable cells behaved similarly to transiently transfected cells in that the double knockdown of Mdm2 and Spry4 was able to rescue migration decreased by Mdm2 silencing alone (Supplementary Fig. 6E). These experiments served as a prelude to the lateral-tail vein injections that was performed as described in Fig 2I. Remarkably, using the same assay our results showed that *in vivo* the silencing of Spry4 is able to completely rescue the effects of Mdm2 loss in metastasis formation *in vivo* (Fig. 6J). Controls with siRNAs and stable shRNAs against Spry4 alone were performed and are shown in Supplementary Figure 6F-K and are reviewed in the Discussion section.

Taken together our findings point to the regulation of Spry4 by Mdm2 as a critical molecular mechanism by which Mdm2 knockdown leads to impaired cell attachment, cell migration, FA formation and development of metastasis *in vivo* in HT1080 cells. Further, Spry4 regulation of cofilin and not ERK activation is the likely mechanism by which Mdm2 regulates those cellular processes.

## Discussion

Mdm2 regulates several activities involved in tumorigenesis, one of which is migration of cancer cells. We bring several new findings concerning the mechanism by which Mdm2 regulates migration in cells that do not express p53.

First, our data strongly support the likelihood that the E3 ligase activity of the Mdm2/MdmX complex is involved in the cellular processes we have studied. Second, Mdm2 is required for robust FA formation in order to maintain the attachment of cells to the ECM and this likely does not involve regulation of EMT. Third, we find that Mdm2 suppression of Spry4 in the cells that we have studied is critical for Mdm2 regulation of migration, FA formation and metastasis.

### The E3 ligase activity of the Mdm2/MdmX complex is required for Mdm2 to promote migration and invasion

To date those studies reporting regulation of Mdm2 altering cell migration have modulated Mdm2 protein levels, either by overexpression of exogenous Mdm2, by silencing of the protein using short hairpin or small interference RNAs or by using inhibitors of Mdm2 such as SP141^50^ and InulanolideA^51^, that reduce Mdm2 protein levels. Although it is clear that modulation of Mdm2 protein levels leads to changes in cell migration, information as to whether the E3-ligase function of the Mdm2/MdmX complex is required for this process is lacking. MEL23, an inhibitor of the Mdm2-MdmX complex E3 ligase activity leads to accumulation of Mdm2 and MdmX ^30^ and treatment with this inhibitor recapitulates all the effects observed in response to Mdm2 silencing.

We showed that MdmX silencing reproduces the effects of Mdm2 downregulation; however, the effects of MdmX silencing on Mdm2 levels vary from cell line to cell line ^52^ and silencing of MdmX in HT1080 cells leads to downregulation of Mdm2 ^53^. This needs to be taken in consideration when analyzing the role of MdmX in the phenotype. Because MdmX levels also decreased Mdm2 protein we cannot exclude the possibility that part of the effect observed in siMdmX conditions is due to Mdm2 decrease and not only because of MdmX knockdown. Nevertheless, our results together strongly point to the involvement of the E3-ligase activity in the regulation of cell migration and related phenotypes induced by Mdm2.

### Mdm2 is required for focal adhesion formation in HT1080 cells

Focal adhesions are essential structures involved in cell anchorage that connect the cell cytoskeleton with membrane receptors, which are in contact with the extracellular environment. As such their function is intrinsic to cell invasion and metastasis. Formation of FAs has been shown to regulate metastasis in different models ^54–57^ where activation of the pathway is associated with increased invasion. In our work we show that the regulation of this process by Mdm2 is needed for cell attachment, migration and metastasis. Loss or inhibition of Mdm2 sets up a unique condition where Spry4 is needed to prevent FA formation.

Although in our model EMT/MET was not induced in response to Mdm2 modulation and thus is apparently not involved in the mechanism by which Mdm2 regulates migration, invasion and metastasis, we do not discard the possibility that in other cell types it may be a significant contributing factor. The same applies to FAs, where, although we demonstrate this regulation happens in two different cell types, more studies are needed to determine the generalizability of our findings.

### Mdm2 down-regulation of Sprouty4 is required for Mdm2 to maintain FA formation and support metastasis

We are the first to associate Mdm2 with the regulation of Sprouty proteins, specifically Spry4. Although we show evidence that Spry4 expression is caused by changes in mRNA being the focus of Mdm2 regulation, we could not rule out as well an effect of Mdm2 on Spry4 protein turnover. It is difficult to assess Spry4 protein stability using cyclohexamide in our experimental conditions due to the very short half-life of Mdm2 (∼ two hours). However, our experiments with the proteasome inhibitor MG-132 suggest that Spry4 regulation is not at the protein level. Furthermore, consistent with our findings, iron depletion reportedly increases expression of Spry4 by increasing both transcription and stability of Spry4 mRNA ^58^.

Overexpression of Spry4 leads to decreased cell migration in the majority of cell models tested to date ^44–47, 59–62^ with a few reports showing no significant difference in migration after overexpression ^63, 64^. By contrast, migration in response to Spry4 knockdown was found to have widely varying effects in different studies, ranging from increasing cell migration ^59, 65, 66^, to having no effect on the process ^67^ or even having the opposite effect, where silencing of Spry4 led to decreased migration ^68, 69^. This reinforces the context dependence of Sprouty family members and more specifically, Spry4. In fact, we found that silencing of Spry4 alone led to decreased migration in HT1080 cells (Supplementary Fig 6F and G), which was unexpected when considering that when silenced together with Mdm2 complete migration was restored. Breast cancer cells depleted of Spry4 were reported to increase the metastatic burden in mice ^65^, while we found that shSpry4 stable cell lines barely metastasized to the lungs using our fibrosarcoma model (Supplementary Figure 6H and I). It is notable that while cell migration was decreased in siSpry4-silenced cells, FA formation was not altered (Supplementary Fig 6J), thereby indicating that while siMdm2 alone and siSpry4 alone each leads to reduced migration, they are likely working through different pathways.

Although Sprouty family members are most well understood as negative regulators of receptor tyrosine kinases (RTKs), through Ras/Raf/ERK activation ^42, 66, 70, 71^, these proteins have been shown to play non-canonical functions in cells as well. Spry4, in particular, has been shown to regulate different kinases/kinase-driven pathways including PI3K/Akt ^69^, PLC/PIP_2_ ^72^, c-Src ^45^ and TESK1 ^48^ independently of their activity on the Ras/Raf/ERK pathway. We demonstrate here that in HT1080 cells, changes in Spry4 do not lead to modulation of ERK phosphorylation (Supplementary Figure 6K), pointing to a non-canonical Spry4 function in regulating migration in this context. We show that phosphorylation of cofilin-1 is involved, at least in part, on the regulation of migration by Mdm2 through Spry4. Although the role of Spry4 in migration and metastasis inhibition induced by reducing Mdm2 levels is clear, further studies are needed to determine the precise mechanism activated by Spry4 silencing alone to induce those changes.

### Mdm2 and promotion of tumor metastasis

Mdm2’s role in primary tumor growth kinetics is still somewhat controversial, with reports of Mdm2 inhibition leading to decreased tumor mass in one study ^18^ and data showing no difference in tumor volume after Mdm2 silencing in another ^22^. Hauck and colleagues showed that silencing of Mdm2 leads to decreased tumor volume in MDA-MB-231 cells but the tumor growth using an MDA-MB-231-derived cell line, TMD231, is not impacted by Mdm2 knockdown ^17^. Its role in metastasis, however, is more consistent with reports of decreased circulating tumor cells as well as decreased metastatic burden when Mdm2 is decreased ^17, 18, 22, 23^.

We found that Mdm2 knockdown affects tumor growth *in vivo* (Supplementary Fig 2D), which might make it difficult to uncouple the roles of Mdm2 in cell migration, attachment, and metastasis alone. However, considering the body of evidence we collected in 2D and 3D culture settings (in which proliferation is not significantly affected), as well as the *in vivo* metastasis models, we suggest that modulation of Mdm2 levels affects the metastatic capabilities independent of its role in tumor cell proliferation and primary tumor growth. This is supported by Gao and colleagues who reported differences in metastatic burden when modulating Mdm2 while tumor growth is unaffected ^22^.

Our work poses several questions. How does Sprouty4 regulate migration without preventing MAPK signaling? Does Mdm2 regulate expression of other Spred-Sprouty family members and is such regulation required for Mdm2 roles in migration and metastasis in different contexts? Finally, how can our findings be exploited for clinical benefit? Our findings reveal the potential of targeting Mdm2 for cancer treatment with a special gain for some patients in advanced stages of the disease. While most inhibitors of Mdm2 in clinical trials target tumors with wild-type p53 with drugs that dissociate Mdm2 and p53, we suggest that blocking the E3 ligase function of Mdm2 might have more general therapeutic benefits.

## Acknowledgments

We are grateful to members of the Prives laboratory for helpful suggestions and discussion. This study was supported by grants CA87497 and CA220526 to CP, as well by grants P30 CA013696, PO1 CA098101 to AR and F31 CA275369 to GE This work was supported by grants from the Japan Society for the Promotion of Science KAKENHI [Grants-in-Aid for Scientific Research (B) #19H03708, #21H02974, (C) #18K07439, (Challenging Exploratory Research) #21K19398]. This work was partly supported by the Uehara Memorial Foundation, Naito Foundation, Princes Takamatsu Cancer Research Fund, Takeda Science Foundation, Yamaguchi Endocrine Research Foundation, Novartis Foundation (Japan) for Promotion of Science, Kose Cosmetology Research Foundation and Medical Institute of Bioregulation Kyushu University Cooperative Research Project Program.

## Authors contribution

RMdQ conceived the idea for the project. RMdQ, AG, GE, AKR and CP designed the experiments. RMdQ, AG, GE, NH and YK conducted the experiments. RMdQ, GE and AG analyzed the results. TT deposited proteomics database. RMdQ wrote the paper with support of CP, AG, GE, NH, TT and CP contributed to manuscript preparation. All authors reviewed the results and approved the final version of the manuscript.

## Material and methods

### Chemicals and Reagents

#### Antibodies

Primary and secondary antibodies used for immunoblotting were at 1:1,000 and 1:2,000 dilution, respectively. Antibodies against Mdm2 (cat.#86934), N-cadherin (cat.#13116S), Tensin-2 (cat.#11990), Talin-1 (cat.#4021), Vinculin (cat.#4650), Paxilin (cat.#12065), alpha-actinin (cat.#6487), RhoB (cat.#2098S), RhoC (cat.#3430S), focal adhesion kinase (FAK) (cat.#3285), integrin beta1 (cat#9699), cofilin-1 (cat.#5175T) and phospho cofilin-1 (Ser3) (cat.#3313T) were purchased from Cell Signaling Technology. Anti-β-actin (A2066 and A2228), anti-fibronectin (cat.#F3648), anti-mouse peroxidase (cat.#A4416) and anti-rabbit peroxidase (cat.#A6154) were purchased from Sigma-Aldrich. Anti-E-cadherin (cat.#sc-8426) and anti-RhoA (cat.#sc-418) were purchased from Santa Cruz Biotechnology. Anti-p-FAK(Tyr397) (cat.#05-1140) was purchased from Millipore. Anti-laminin (cat.#AHP2491) was purchased from Bio-Rad. Anti-α-tubulin (cat.#AA4.3) was purchased from DSHB. Anti-sprouty4 (cat.#A04343-2), anti-integrin alpha-3 (cat#A02902) and anti-integrin alpha-2 (cat#A01933-2) were purchased from Boster Biological Technology. Anti-vinculin AlexaFluor647 (cat.#ab196579) was purchased from Abcam. Phalloidin AlexaFluor Plus 555 (cat.#A30106) was purchased from Thermo Fisher Scientific. Anti-p53 DO-I and 1801 were purified from hybridomas produced in-house. Anti-MdmX mAb 8C6 was produced in Dr. Jiandong Chen’s lab and kindly gifted to our group. **Constructs:** For Mdm2 transient knockdown cells were transfected with Silencer Select siRNA s8630 (siMdm2#1) or s224037 (siMdm2#2). For MdmX knockdown cells were transfected with FlexiTube siRNA Mdm4 (cat.#SI00037163, siMdmX#4) from QIAGEN or costume siRNA against MdmX (siMdmX#1, sequence: 5’-AGAUUCAGCUGGUUAUUAA-3’) from Sigma-Aldrich. For Spry4 knockdown cells were transfected with a pool of 3 siRNAs against Spry4 (s37824, s37825, s37826). Silencer Select Negative Control siRNA #1 (cat.#4390843) was used as transfection control. All siRNAs but the ones against MdmX were purchased from Ambion and transfection was done using Lipofectamine RNAiMax transfection reagent (cat.#13778150, Thermo Scientific). Stable cell lines were stablished transducing cells with a set of 4 shRNAs against Mdm2 (Origene, cat.#TL311529) or shRNA negative control (Origene, cat.#TR30021) and using the adequate drug for cell selection. For shSpry4 stable cell lines, cells were transduced with a set of 4 shRNAs against Spry4 (OriGene, cat.#HC108594) or shRNA negative control (OriGene, cat.#TR30033), and using the appropriate drug for cell selection.

#### Drugs and other reagents

MEL23 (Sigma-Aldrich, cat#373227), MG132 (Selleck Chemicals, cat.#S2619), Accutase (Thermo Fisher Scientific, cat.#ICN1000449), Actinomycin D (Cayman Chemichals, cat#11421), DRB (Sigma-Aldrich, cat#D1916), CellTracker dye (Thermo Fisher Scientific, cat.#C34551), pepsin-treated (PT) bovine type I collagen (Advanced BioMatrix, cat.#5010), growth factor-reduced Phenol Red-free BME/Matrigel (8.9-10 mg/ml solution from BD Biosciences), DMEM solution (10×), NaOH (1 N) and sodium bicarbonate solution (7.5%) (Sigma Aldrich), 4-(2-hydroxyethyl)-1-piperazineethanesulfonic acid (HEPES) buffer (1 M) (Invitrogen, cat.#15630080), CellMask membrane dye (Thermo Fisher Scientific, cat.#C10045), Alexa Fluor Plus 555 Phalloidin (Thermo Fisher Scientific, cat.#A30106), Molecular Probes Deoxyribonuclease I Alexa Fluor 488 Conjugate (Thermo Fisher Scientific, cat.#D12371).

### Cell culture

Human fibrosarcoma cell line CRISPR-engineered HT1080 p53 KO cells ^53^ were grown in Dulbecco’s modified Eagle’s Medium (DMEM) (cat.#12100-061, Gibco) supplemented with 10% heat-inactivated fetal bovine serum (FBS) (cat.#900108H, Gemini), 100 units of penicillin, and 100 mg/ml streptomycin (cat.#15140122, Thermo Scientific). Human lung adenocarcinoma H1299 cells were grown in RPMI 1640 medium (cat.#11875119, Thermo Scienftific) supplemented with 10% heat-inactivated FBS, 100 units of penicillin, and 100 mg/ml streptomycin. Cells were maintained at 37°C with 5% CO_2_ and were sub-cultured using trypsin-EDTA (cat.#15090-046, Gibco) every 2 or 3 days. All cells used in this work were previously tested for mycoplasma contamination and found to be negative.

### Mdm2 and Mdm2/Spry4 knockdown stable cell lines

To perform *in vivo* experiments we stablished a Mdm2 knockdown stable cell line and a Mdm2/Sprouty4 double knockdown stable cell line, as well as control cell lines expressing shScramble contructs. For the single knockdown, HT1080 p53KO cells were transduced with lentivirus carrying a pool of shRNAs against Mdm2 (4 different shRNAs purchased from OriGene, cat.#TL311529) or shScramble sequence as control (OriGene, cat.#TR30021). Constructs co-expressed GFP. Stable cell lines were generated by selecting transduced cells with 1 µg/ml puromycin for 72 h. Cells were kept on media with a lower concentration (0.5 µg/ml) of puromycin after selection for one week then challenged with 1 µg/ml puromycin every 4 weeks in culture.

Cells expressing shMdm2 or shScramble were sorted for GFP positive cells and transduced with lentivirus carrying a pool of shRNAs against Sprouty4 (4 different shRNAs purchased from OriGene, cat.#HC108594) or shScramble sequence as control (OriGene, cat.#TR30033). Constructs co-expressed RFP. Stable cell lines were generated by selecting transduced cells with 5 µg/ml blasticidin for 72 h. Cells were kept on media with lower concentration (2 µg/ml) of blasticidin after selection for one week then challenged with 5 µg/ml blasticidin every 4 weeks in culture. Newly established cell lines were sorted for GFP- and RFP-positive cells.

### Cell viability

To ensure the treatments used in this work did not interfere with cell viability cells were stained with Trypan Blue (BioRad, cat.#1450011) 48 h after transfections with siRNA or after 24 h of treatment with MEL23 or vehicle. Cell viability in 3D was determined using CellTiter Glo (Promega, cat.#G9681) according to manufacturer’s instructions. Luminescence was measured by a BioTek SynergyH1 Hybrid Reader.

### Immunoblotting

After desired treatments, cells were harvested with trypsin, pelleted and lysed in RIPA buffer (50 mM Tris, pH 8, 0.5% deoxycholate, 0.1% SDS, 1% Nonidet P-40, 5 mM EDTA, 150 mM NaCl) along with protease and phosphatase inhibitors. Samples were loaded in equal protein amounts and polypeptides were separated by SDS-PAGE. Proteins were transferred to a PVDF membrane (IPVH00010, Immobilon, Millipore) and blocked for 1 h with Tris-buffered saline containing 0.05% Tween 20 and 3% bovine serum albumin. Membranes were incubated overnight at 4 °C with primary antibody followed by 3 washes with Tris-buffered saline, 0.05% Tween 20. Secondary HRP-conjugated antibody was added and incubated for 1 h at room temperature (RT). After washes, blots were visualized by chemiluminescence detection using chemiluminescent horseradish peroxidase reagents (Thermo Fisher, Pierce, cat# 32106 or EMD Millipore, Immobilon, cat# WBKLS0050) according to the manufacturer’s instructions. Quantification of protein expression was performed using ImageJ software.

### Co-immunoprecipitation

Cells were seeded in 10 cm dishes, and after desired treatments MG132 was added for 4 hours, followed by lysing samples in NP-40 lysis buffer. Lysates (400 µg of protein) were then incubated with primary antibody or IgG control antibody for 2 h at 4 °C with constant rotation, followed by 1 h incubation with Protein G agarose beads (Millipore, catalog no. 16-266) at 4 °C with constant rotation. Beads were washed 4 times with lysis buffer or PBS then protein was eluted from beads with Laemmli buffer and boiled for 5 min. Samples were analyzed by immunoblotting.

### Quantitative Real Time PCR (qPCR)

Total RNA was isolated using a Qiagen RNeasy Minikit (Qiagen, cat.#74106) according to the manufacturer’s instructions. A total of 1 μg of RNA was used for reverse transcription (RT) using QuantiTect Reverse Transcription Kit (cat.#205311). The cDNA products were diluted (1:10) with nuclease-free water and analyzed by qPCR using Power SYBR Green PCR Master Mix (Applied Biosystems, cat.#4368708) according to the manufacturer’s instructions. Quantitative PCR analysis was performed using the following protocol: polymerase activation and DNA denaturation for 30 s at 95 °C; amplification denaturation for 5 s at 95 °C and annealing for 30 s at 60 or 62 °C with 40 cycles; and melt curve 65–95 °C with 0.5 °C increment 5 s/step. Quantification cycle (Cq) value was recorded by StepOne software. Relative changes in cDNA levels were calculated using the comparative Ct method (ΔΔCT method). Transcripts were normalized to RPL32. Primers for SPRY4: forward 5’- AGAAGTGTACTGAAGGGACTGGAG-3’; reverse 5’- GTGTGTAGACCACCAAGATCACC-3’. Primers for MDM2: forward 5’- TTGGCGTGCCAAGCTTCTCT-3’, reverse 5’-TACCTGAGTCCGATGATTCC-3’. Primers for ARG1: forward 5’-GTCTGTGGGAAAAGCAAGCG-3’, reverse 5’- CACCAGGCTGATTCTTCCGT-3’. Primers for RPL32: forward 5’- TTCCTGGTCCACAACGTCAAG-3’; reverse 5’-TGTGAGCGATCTCGGCAC-3’.

### Immunofluorescence

Cells were seeded in 6-well plates and after desired treatments, were detached using PBS-EDTA 5mM and reseeded onto coverslips. For cell spreading analysis and focal adhesion staining, cells were fixed 6 h after seeded in collagen-I coated (Advanced BioMatrix, cat.#5056) coverslips. Once ready to be processed cells were fixed and permeabilized in one step with paraformaldehyde (PFA) 4% - 0.2% Triton X-100 for 10 minutes at room temperature. After washes with PBS cells were blocked in PBS-1%-BSA for 30 min and incubated with anti-vinculin antibody overnight in a humid chamber. Coverslips were then washed with PBS and incubated with fluorescent phalloidin for 1 hour RT, washed with PBS and incubated with DAPI (5 µM) for 10 minutes. Coverslips were mounted and micrographs were acquired using ZEISS LSM800. Quantifications were performed using Fiji software.

### F/G actin staining

Filamented actin (F-actin) was stained using fluorescently-congujated phalloidin while monomeric actin (G-actin) was probed using fluorescently-labeled DNase I. Cells were fixed in PFA 4% for 10 minutes, washed with PBS and permeabilized with PBS-0.12% Triton X-100 for 5 minutes RT. After washes with PBS cells were incubated with fluorescent phalloidin and DNAse I in manufecturer’s suggested concentrations for 1 hour RT, washed with PBS and incubated with DAPI for 10 minutes. Coverslips were mounted and micrographs were acquired using ZEISS LSM800.

### Cell motility/migration assay

Cell migration was measured by the wound scratch assay. Briefly, cells were seeded in 6-well plates and allowed to reach 100% confluence. The confluent layers were each subjected to a scratch with a 200 µl tip, then washed and media added back. HT1080 cells were serum starved and treated with 2 µg/ml mitocycin-C, while H1299 cells were kept in 2% FBS-RPMI with 2 µg/ml mitocycin-C for the entire duration of the experiment. At the time of scratching, 0 h micrographs were taken and after the indicated time intervals, micrographs of the same points in the wound were taken for comparison. Quantification of wound area was performed using ImageJ software, time. At 0 h the images were considered to have 100% wound areas.

### Determination of cell size in solution

To analyze the cell size in solution cells (2.5 x 10^5^) were seeded in 6-well plates. After treatments cells were detached and cell size was analyzed by forward scatter (FSC) by flow cytometry in a FACS Celesta, using FACS diva software (BD).

### ECM attachment assay

Cells were seeded in 6-well plates, after desired treatments cells were detached using PBS-EDTA 5mM and collected. After washing and resuspension in FBS free media, 10^5^ cells were reseeded on Laminin-(Corning, cat.#354412), Fibronectin-(Corning, cat.#356271), Collagen I-coated (Corning, cat.#356271) 24-well, or on uncoated 24-well plates, all of which had been pre blocked with PBS 1% BSA. Plates were kept at 37 °C with 5% CO_2_ for 1 h. After incubation the culture dish wells were washed three times with FBS-free media to remove unattached cells. The plates were then kept in complete media for 1 h at 37 °C with 5% CO_2_ for cell recovery and then remaining cells were stained using 0.2% crystal violet and 10% ethanol. Micrographs of crystal violet-stained wells were taken and followed by dissolving material on the wells in 2% SDS for quantification. Absorbance was measured at 540 nm by BioTek SynergyH1 Hybrid Reader and was normalized to control conditions.

### Analysis of mRNA stability

mRNA stability was assessed by treatment with two transcription inhibitors: actinomycin D (actD) as described previously ^73^ or 5, 6-dichloro-1-β-D-ribofuranosylbenzimidazole (DRB)^74^. Briefly, cells were seeded in 12-well plates and treated with MEL23 for 24 h. Then, 10 µg/ml of act D or 100 µM of DRB were added to wells for indicated time intervals. Cells were harvested, RNA was isolated and cDNA was generated as described above. mRNA levels were analyzed by qPCR. Data of actD-or DRB-treated cells were normalized to cells treated with vehicle in each condition (DMSO or MEL23 treatment).

### Luciferase assay

A Spry4 Firefly luciferase reporter plasmid containing the promoter region of Spry4 (kindly provided by Dr. Meredith Tennis, University of Colorado, USA)^46^ or a negative control were used in co-transfection assays with Renilla-luciferase expressing plasmids, as internal controls. Briefly, cells were seeded in 12-well plates and transfected the next day with siRNA directed against Mdm2. After 24 h cells were co-transfected with 3.9 µg of a Firefly luciferase-expressing construct (Spry4 promoter or mock) and 0.1 µg of Renilla-luciferase expressing plasmid. Firefly and Renilla signals were measured 24 h after transfection using the Dual-Glo Luciferase Assay System (Promega, cat#E2920) according to manufacturer’s instructions. Luminescence was measured by BioTek SynergyH1 Hybrid Reader and the Firefly signal was normalized to the Renilla signal.

### Untargeted proteomics

Cells were transfected using siRNA control or 2 different siRNAs against Mdm2 for 24 h for siRNA sample set; or treated with MEL23 (7 µM) or DMSO for 24 h for drug sample set. Then, proteins were isolated from the cell samples using a phenol-guanidinium isothiocyanate (P/GTC) reagent according to the manufacturer’s protocol ^75, 76^. After RNA was precipitated from the aqueous phase with isopropanol, proteins were precipitated from the phenol/ethanol phase by the addition of acetone. The protein precipitate was redissolved in 2% SDS and 100-mM Tris-HCl (pH 8.5) using a sonicator (BIORUPTOR II, CosmoBio, Tokyo, Japan). The protein extract was measured using a BCA Protein Assay Kit (Thermo Fisher Scientific, Waltham, MA, USA) and adjusted to 1 mg/ml with 2% SDS and 100 mM Tris-HCl, pH 8.5. Then 20 µL of protein extract was digested with trypsin and Lys-C, and the peptides were analyzed by LC-MS/MS (Orbitrap Exploris 480 Mass Spectrometer, Thermo Fisher Scientific) as previously reported ^77^. The MS files were searched against an in silico human spectral library using Scaffold DIA (Proteome Software, Inc., Portland, OR, USA) as previously reported ^78^. The mass spectrometry proteomics data have been deposited in the ProteomeXchange Consortium (http://proteomecentral.proteomexchange.org) via the jPOST partner repository (http://jpostdb.org) with the dataset identifier PXD033789.

For the proteomics analysis, volcano plots were generated using VolcaNoseR ^79^. Heatmaps showing significantly differentially expressed proteins were generated using Heatmapper ^80^. Pathway analysis was performed using Enrichr ^25, 81, 82^.

### Database analysis of gene expression and protein interaction

Analysis of Spry4 gene expression in normal tissue and tumors was performed using GEPIA ^83^, grouping patient samples from the TCGA and GTEx projects.

### Invasion assay and 3D morphology in hydrogels

<u>Generation and treatment of multicellular tumor spheroids:</u> Cells were seeded in 6-well plates and either silenced for Mdm2 using different siRNAs or treated with MEL23 (7 µM) or DMSO for 24 h. After desired treatments cells were detached using Accutase. Spheroids were formed in appropriate growth medium supplemented with 0.25 mg/ml BME (Basement Membrane Extract) using a centrifugation method described previously ^84^. Spheroids were allowed to form for 24 h at 37°C under 5% carbon dioxide. For MEL23 treatment, the same concentration of inhibitor was added to the biomatrix and the overlay media to maintain continuous treatment during the invasion assay. <u>Preparation of collagen solutions</u>: collagen (1 mg/ml) and collagen/BME matrices were prepared as described in Guzman et al, Biomaterials 2014 ^35^. <u>Preparation of cell-embedded gels</u>: to prepare gels loaded with a single spheroid, the collagen solution was added to a chamber and the spheroid was added to the liquid collagen which was then transferred to a 37°C incubator for gelation. For collagen gels loaded with dispersed cells, the collagen solution was prepared omitting 50% of the water and neutralized. The water volume was substituted with growth medium containing the desired number of cells (2 X 104 – 4 X 104 cells/ml). Subsequently cell-loaded collagen was added to the chamber and gelled at 37°C. <u>Immunocytochemistry and labeling</u>: For staining the actin cytoskeleton in collagen-embedded cells samples were fixed with 4% PFA 15 minutes at RT, permeabilized with 0.2% TritonX/H2O for 10 min at RT and incubated with fluorescently labeled phalloidin dye for 16-20 h at 4°C. <u>Spheroid invasion</u>: for invasion analysis, spheroids were imaged at 1 h and 18 h after implantation. For spheroids invading in pure collagen I gels, magnified fluorescence images were used to determine the invaded area, defined as the difference between the area of the spheroid at t=1h and an ellipse that circumscribes 90% of the invasive cells at t=18 h. For spheroids invading in collagen/BME composite matrices, maximum intensity projections were thresholded and used to generate masks from which the non-invading spherical core of the spheroid was removed. The remaining invasive structures were analyzed. The total area of invasive structures per spheroid was used to assess invasion efficiency.

### Lateral tail-vein injections and orthotopic implantation

All animal studies were approved by the Columbia University Institutional Animal Care and Use Committee. To establish the lung metastasis model, 5-week-old male and female athymic nude mice were purchased (Taconic). At the age of 7 weeks, 1.0×10^6^ (tail-vein) and 5.0×10^5^ (orthotopics) cells were resuspended in 100 μL of cold Dulbecco’s phosphate-buffered saline (DPBS) and injected into the distal end of the lateral tail vein (tail-vein) or right quadriceps femoris muscle (orthotopics) with a 28-gauge 0.5 in needle. After injection, mice were monitored daily for signs of pain or discomfort and weight loss. To detect the formation of lung metastatic lesions, mice were euthanized at 8-weeks (tail-vein) or 6-weeks (ortothopics) time points by CO_2_ and cervical dislocation. To identify lung metastases, the mouse lungs were flushed with 20 mL Heparin-PBS to deplete red blood cells and were imaged with Keyence fluorescent microscope at 2X magnification.

### Statistical Analysis

Data reported in this paper were expressed as the mean ± S.D. from at least three independent experiments. A significant difference from the respective control for each experimental test condition was assessed by one or two-way analysis of variance with respective post-tests, or Student’s t test using GraphPad Prism 7 software. Values of p < 0.05 were considered statistically significant.

## References

1. Fakharzadeh, S.S., Trusko, S.P. & George, D.L. Tumorigenic potential associated with enhanced expression of a gene that is amplified in a mouse tumor cell line. EMBO J 10, 1565–1569 (1991).

2. Karni-Schmidt, O., Lokshin, M. & Prives, C. The Roles of MDM2 and MDMX in Cancer. Annu Rev Pathol 11, 617–644 (2016).

3. Iwakuma, T. & Lozano, G. MDM2, an introduction. Mol Cancer Res 1, 993–1000 (2003).

4. Manfredi, J.J. The Mdm2-p53 relationship evolves: Mdm2 swings both ways as an oncogene and a tumor suppressor. Genes Dev 24, 1580–1589 (2010).

5. Tackmann, N.R. & Zhang, Y. Mouse modelling of the MDM2/MDMX-p53 signalling axis. J Mol Cell Biol 9, 34–44 (2017).

6. Klein, A.M., de Queiroz, R.M., Venkatesh, D. & Prives, C. The roles and regulation of MDM2 and MDMX: it is not just about p53. Genes Dev 35, 575–601 (2021).

7. Donehower, L.A. & Lozano, G. 20 years studying p53 functions in genetically engineered mice. Nat Rev Cancer 9, 831–841 (2009).

8. Bagci, O. & Kurtgoz, S. Amplification of Cellular Oncogenes in Solid Tumors. N Am J Med Sci 7, 341–346 (2015).

9. Berberich, S.J. Mdm2 and MdmX involvement in human cancer. Subcell Biochem 85, 263–280 (2014).

10. Hou, H., Sun, D. & Zhang, X. The role of MDM2 amplification and overexpression in therapeutic resistance of malignant tumors. Cancer Cell Int 19, 216 (2019).

11. Eischen, C.M. Role of Mdm2 and Mdmx in DNA repair. J Mol Cell Biol 9, 69–73 (2017).

12. Wienken, M., Moll, U.M. & Dobbelstein, M. Mdm2 as a chromatin modifier. J Mol Cell Biol 9, 74–80 (2017).

13. Feeley, K.P., Adams, C.M., Mitra, R. & Eischen, C.M. Mdm2 Is Required for Survival and Growth of p53-Deficient Cancer Cells. Cancer Res 77, 3823–3833 (2017).

14. Klein, A.M. et al. MDM2, MDMX, and p73 regulate cell-cycle progression in the absence of wild-type p53. Proc Natl Acad Sci U S A 118 (2021).

15. Miao, Z., Liu, S., Xiao, X. & Li, D. LINC00342 regulates cell proliferation, apoptosis, migration and invasion in colon adenocarcinoma via miR-545-5p/MDM2 axis. Gene 743, 144604 (2020).

16. Sha, M.X., Huang, X.W. & Yin, Q. MiR-548b-3p inhibits proliferation and migration of breast cancer cells by targeting MDM2. Eur Rev Med Pharmacol Sci 24, 3105–3112 (2020).

17. Hauck, P.M. et al. Early-Stage Metastasis Requires Mdm2 and Not p53 Gain of Function. Mol Cancer Res 15, 1598–1607 (2017).

18. Wang, W. et al. A novel inhibitor of MDM2 oncogene blocks metastasis of hepatocellular carcinoma and overcomes chemoresistance. Genes Dis 6, 419–430 (2019).

19. Jana, A. et al. NFkB is essential for activin-induced colorectal cancer migration via upregulation of PI3K-MDM2 pathway. Oncotarget 8, 37377–37393 (2017).

20. Jiang, D. et al. MiR-758-3p suppresses proliferation, migration and invasion of hepatocellular carcinoma cells via targeting MDM2 and mTOR. Biomed Pharmacother 96, 535–544 (2017).

21. Qiu, W., Xia, X., Qiu, Z., Guo, M. & Yang, Z. RasGRP3 controls cell proliferation and migration in papillary thyroid cancer by regulating the Akt-MDM2 pathway. Gene 633, 35–41 (2017).

22. Gao, C. et al. Context-dependent roles of MDMX (MDM4) and MDM2 in breast cancer proliferation and circulating tumor cells. Breast Cancer Res 21, 5 (2019).

23. Chen, Y. et al. MDM2 promotes epithelial-mesenchymal transition and metastasis of ovarian cancer SKOV3 cells. Br J Cancer 117, 1192–1201 (2017).

24. Bradbury, R., Jiang, W.G. & Cui, Y.X. MDM2 and PSMA Play Inhibitory Roles in Metastatic Breast Cancer Cells Through Regulation of Matrix Metalloproteinases. Anticancer Res 36, 1143–1151 (2016).

25. Chen, E.Y. et al. Enrichr: interactive and collaborative HTML5 gene list enrichment analysis tool. BMC Bioinformatics 14, 128 (2013).

26. Yan, Y., Peng, Y., Ou, Y. & Jiang, Y. MicroRNA-610 is downregulated in glioma cells, and inhibits proliferation and motility by directly targeting MDM2. Mol Med Rep 14, 2657–2664 (2016).

27. Wu, D. et al. MicroRNA-379-5p plays a tumor-suppressive role in human bladder cancer growth and metastasis by directly targeting MDM2. Oncol Rep 37, 3502–3508 (2017).

28. Liu, L. et al. CP31398 attenuates endometrial cancer cell invasion, metastasis and resistance to apoptosis by downregulating MDM2 expression. Int J Oncol 54, 942–954 (2019).

29. Polanski, R. et al. MDM2 interacts with NME2 (non-metastatic cells 2, protein) and suppresses the ability of NME2 to negatively regulate cell motility. Carcinogenesis 32, 1133–1142 (2011).

30. Herman, A.G. et al. Discovery of Mdm2-MdmX E3 ligase inhibitors using a cell-based ubiquitination assay. Cancer Discov 1, 312–325 (2011).

31. Qiang, P., Shao, Y., Sun, Y.P., Zhang, J. & Chen, L.J. Metformin inhibits proliferation and migration of endometrial cancer cells through regulating PI3K/AKT/MDM2 pathway. Eur Rev Med Pharmacol Sci 23, 1778–1785 (2019).

32. Yang, J.Y. et al. MDM2 promotes cell motility and invasiveness by regulating E-cadherin degradation. Mol Cell Biol 26, 7269–7282 (2006).

33. Galarza, S., Kim, H., Atay, N., Peyton, S.R. & Munson, J.M. 2D or 3D? How cell motility measurements are conserved across dimensions in vitro and translate in vivo. Bioeng Transl Med 5, e10148 (2020).

34. Baskaran, J.P. et al. Cell shape, and not 2D migration, predicts extracellular matrix-driven 3D cell invasion in breast cancer. APL Bioeng 4, 026105 (2020).

35. Guzman, A., Ziperstein, M.J. & Kaufman, L.J. The effect of fibrillar matrix architecture on tumor cell invasion of physically challenging environments. Biomaterials 35, 6954–6963 (2014).

36. Poincloux, R., Lizarraga, F. & Chavrier, P. Matrix invasion by tumour cells: a focus on MT1-MMP trafficking to invadopodia. J Cell Sci 122, 3015–3024 (2009).

37. Janiszewska, M., Primi, M.C. & Izard, T. Cell adhesion in cancer: Beyond the migration of single cells. J Biol Chem 295, 2495–2505 (2020).

38. Rikitake, Y. & Takai, Y. Directional cell migration regulation by small G proteins, nectin-like molecule-5, and afadin. Int Rev Cell Mol Biol 287, 97–143 (2011).

39. Burridge, K. & Guilluy, C. Focal adhesions, stress fibers and mechanical tension. Exp Cell Res 343, 14–20 (2016).

40. Henning Stumpf, B., Ambriovic-Ristov, A., Radenovic, A. & Smith, A.S. Recent Advances and Prospects in the Research of Nascent Adhesions. Front Physiol 11, 574371 (2020).

41. Ibata, N. & Terentjev, E.M. Development of Nascent Focal Adhesions in Spreading Cells. Biophys J 119, 2063–2073 (2020).

42. Kawazoe, T. & Taniguchi, K. The Sprouty/Spred family as tumor suppressors: Coming of age. Cancer Sci 110, 1525–1535 (2019).

43. Masoumi-Moghaddam, S., Amini, A. & Morris, D.L. The developing story of Sprouty and cancer. Cancer Metastasis Rev 33, 695–720 (2014).

44. Celik-Selvi, B.E. et al. Sprouty3 and Sprouty4, Two Members of a Family Known to Inhibit FGF-Mediated Signaling, Exert Opposing Roles on Proliferation and Migration of Glioblastoma-Derived Cells. Cells 8 (2019).

45. Gong, Y. et al. Sprouty4 regulates endothelial cell migration via modulating integrin beta3 stability through c-Src. Angiogenesis 16, 861–875 (2013).

46. Tennis, M.A. et al. Sprouty-4 inhibits transformed cell growth, migration and invasion, and epithelial-mesenchymal transition, and is regulated by Wnt7A through PPARgamma in non-small cell lung cancer. Mol Cancer Res 8, 833–843 (2010).

47. Jaggi, F., Cabrita, M.A., Perl, A.K. & Christofori, G. Modulation of endocrine pancreas development but not beta-cell carcinogenesis by Sprouty4. Mol Cancer Res 6, 468–482 (2008).

48. Tsumura, Y., Toshima, J., Leeksma, O.C., Ohashi, K. & Mizuno, K. Sprouty-4 negatively regulates cell spreading by inhibiting the kinase activity of testicular protein kinase. Biochem J 387, 627–637 (2005).

49. Tribollet, V. et al. ERRalpha coordinates actin and focal adhesion dynamics. Cancer Gene Ther (2022).

50. Wang, W. et al. Identification of a new class of MDM2 inhibitor that inhibits growth of orthotopic pancreatic tumors in mice. Gastroenterology 147, 893–902 e892 (2014).

51. Qin, J.J. et al. Inulanolide A as a new dual inhibitor of NFAT1-MDM2 pathway for breast cancer therapy. Oncotarget 7, 32566–32578 (2016).

52. Hu, B., Gilkes, D.M., Farooqi, B., Sebti, S.M. & Chen, J. MDMX overexpression prevents p53 activation by the MDM2 inhibitor Nutlin. J Biol Chem 281, 33030–33035 (2006).

53. Venkatesh, D. et al. MDM2 and MDMX promote ferroptosis by PPARalpha-mediated lipid remodeling. Genes Dev 34, 526–543 (2020).

54. Zhang, P. et al. CPNE8 Promotes Gastric Cancer Metastasis by Modulating Focal Adhesion Pathway and Tumor Microenvironment. Int J Biol Sci 18, 4932–4949 (2022).

55. Sun, T. et al. G9a Promotes Invasion and Metastasis of Non-Small Cell Lung Cancer through Enhancing Focal Adhesion Kinase Activation via NF-kappaB Signaling Pathway. Mol Cancer Res 19, 429–440 (2021).

56. Zhong, Y. et al. MYH9-dependent polarization of ATG9B promotes colorectal cancer metastasis by accelerating focal adhesion assembly. Cell Death Differ 28, 3251–3269 (2021).

57. Shen, J. et al. Hippo component YAP promotes focal adhesion and tumour aggressiveness via transcriptionally activating THBS1/FAK signalling in breast cancer. J Exp Clin Cancer Res 37, 175 (2018).

58. Haigl, B., Mayer, C.E., Siegwart, G. & Sutterluty, H. Sprouty4 levels are increased under hypoxic conditions by enhanced mRNA stability and transcription. Biol Chem 391, 813–821 (2010).

59. Vanas, V., Muhlbacher, E., Kral, R. & Sutterluty-Fall, H. Sprouty4 interferes with cell proliferation and migration of breast cancer-derived cell lines. Tumour Biol 35, 4447–4456 (2014).

60. Wang, J., Thompson, B., Ren, C., Ittmann, M. & Kwabi-Addo, B. Sprouty4, a suppressor of tumor cell motility, is down regulated by DNA methylation in human prostate cancer. Prostate 66, 613–624 (2006).

61. Guo, J. et al. SPRY4 suppresses proliferation and induces apoptosis of colorectal cancer cells by repressing oncogene EZH2. Aging (Albany NY) 13, 11665–11677 (2021).

62. Zhang, E. et al. H3K27 acetylation activated-long non-coding RNA CCAT1 affects cell proliferation and migration by regulating SPRY4 and HOXB13 expression in esophageal squamous cell carcinoma. Nucleic Acids Res 45, 3086–3101 (2017).

63. Rathmanner, N. et al. Sprouty2 but not Sprouty4 is a potent inhibitor of cell proliferation and migration of osteosarcoma cells. FEBS Lett 587, 2597–2605 (2013).

64. Stutz, A., Kamptner, A.Z.M. & Sutterluty, H. A Sprouty4 Mutation Identified in Kallmann Syndrome Increases the Inhibitory Potency of the Protein towards FGF and Connected Processes. Int J Mol Sci 22 (2021).

65. Jing, H. et al. Suppression of Spry4 enhances cancer stem cell properties of human MDA-MB-231 breast carcinoma cells. Cancer Cell Int 16, 19 (2016).

66. Qiu, B. et al. Sprouty4 correlates with favorable prognosis in perihilar cholangiocarcinoma by blocking the FGFR-ERK signaling pathway and arresting the cell cycle. EBioMedicine 50, 166–177 (2019).

67. So, W.K. et al. Sprouty4 mediates amphiregulin-induced down-regulation of E-cadherin and cell invasion in human ovarian cancer cells. Tumour Biol 37, 9197–9207 (2016).

68. Pan, Y. et al. LINC00675 Suppresses Cell Proliferation and Migration via Downregulating the H3K4me2 Level at the SPRY4 Promoter in Gastric Cancer. Mol Ther Nucleic Acids 22, 766–778 (2020).

69. Das, M.K., Furu, K., Evensen, H.F., Haugen, O.P. & Haugen, T.B. Knockdown of SPRY4 and SPRY4-IT1 inhibits cell growth and phosphorylation of Akt in human testicular germ cell tumours. Sci Rep 8, 2462 (2018).

70. Brock, E.J. et al. Sprouty4 negatively regulates ERK/MAPK signaling and the transition from in situ to invasive breast ductal carcinoma. PLoS One 16, e0252314 (2021).

71. Sasaki, A. et al. Mammalian Sprouty4 suppresses Ras-independent ERK activation by binding to Raf1. Nat Cell Biol 5, 427–432 (2003).

72. Ayada, T. et al. Sprouty4 negatively regulates protein kinase C activation by inhibiting phosphatidylinositol 4,5-biphosphate hydrolysis. Oncogene 28, 1076–1088 (2009).

73. Ratnadiwakara, M. & Anko, M.L. mRNA Stability Assay Using transcription inhibition by Actinomycin D in Mouse Pluripotent Stem Cells. Bio Protoc 8, e3072 (2018).

74. Radha, K.S. et al. Iron-mediated stability of PAI-1 mRNA in adenocarcinoma cells-involvement of a mRNA-binding nuclear protein. Thromb Res 116, 255–263 (2005).

75. Kawashima, Y. et al. Proteogenomic Analyses of Cellular Lysates Using a Phenol-Guanidinium Thiocyanate Reagent. J Proteome Res 18, 301–308 (2019).

76. Ochiiwa, H. et al. TAS4464, a NEDD8-activating enzyme inhibitor, activates both intrinsic and extrinsic apoptotic pathways via c-Myc-mediated regulation in acute myeloid leukemia. Oncogene 40, 1217–1230 (2021).

77. Nakamura, R. et al. A Simple Method for In-Depth Proteome Analysis of Mammalian Cell Culture Conditioned Media Containing Fetal Bovine Serum. Int J Mol Sci 22 (2021).

78. Takemori, A., Kawashima, Y. & Takemori, N. Bottom-up/cross-linking mass spectrometry via simplified sample processing on anion-exchange solid-phase extraction spin column. Chem Commun (Camb) 58, 775–778 (2022).

79. Goedhart, J. & Luijsterburg, M.S. VolcaNoseR is a web app for creating, exploring, labeling and sharing volcano plots. Sci Rep 10, 20560 (2020).

80. Babicki, S. et al. Heatmapper: web-enabled heat mapping for all. Nucleic Acids Res 44, W147–153 (2016).

81. Xie, Z. et al. Gene Set Knowledge Discovery with Enrichr. Curr Protoc 1, e90 (2021).

82. Kuleshov, M.V. et al. Enrichr: a comprehensive gene set enrichment analysis web server 2016 update. Nucleic Acids Res 44, W90–97 (2016).

83. Tang, Z. et al. GEPIA: a web server for cancer and normal gene expression profiling and interactive analyses. Nucleic Acids Res 45, W98–W102 (2017).

84. Ivascu, A. & Kubbies, M. Rapid generation of single-tumor spheroids for high-throughput cell function and toxicity analysis. J Biomol Screen 11, 922–932 (2006).

