## Supplementary tables and figures for "Sprouty4 is required for Mdm2 regulation of invasion, focal adhesion formation and metastasis in cells lacking p53"

List of content:

Supplementary Table 1 ……………………………………………………….page 2

**Supplementary Table 1:** Integrin expression in HT1080 p53KO cells in response to Mdm2 silencing or treatment with MEL23 for 24 h analyzed by mass spectrometry. Experiment was performed in 3 biological replicates, *p<0.05, **p<0.01, NS: not significant; down: downregulated; up: upregulated.

| | siRNAs | MEL23 |
|---|---|---|
| Integrin alpha 1 | NS | NS |
| Integrin alpha 2 | NS | NS |
| Integrin alpha 3 | NS | *, down |
| Integrin alpha 5 | NS | NS |
| Integrin alpha 6 | NS | *, down |
| Integrin alpha 10 | NS | **, down |
| Integrin alpha V | NS | NS |
| Integrin beta 1 | NS | NS |
| Integrin beta 2 | NS | NS |
| Integrin beta 3 | NS | NS |
| Integrin beta 5 | NS | *, down |
| Integrin beta 8 | NS | NS |
| Integrin beta like 1 | *, up with si#2 | *, down |
| Integrin beta 1 binging protein 1 | NS | NS |

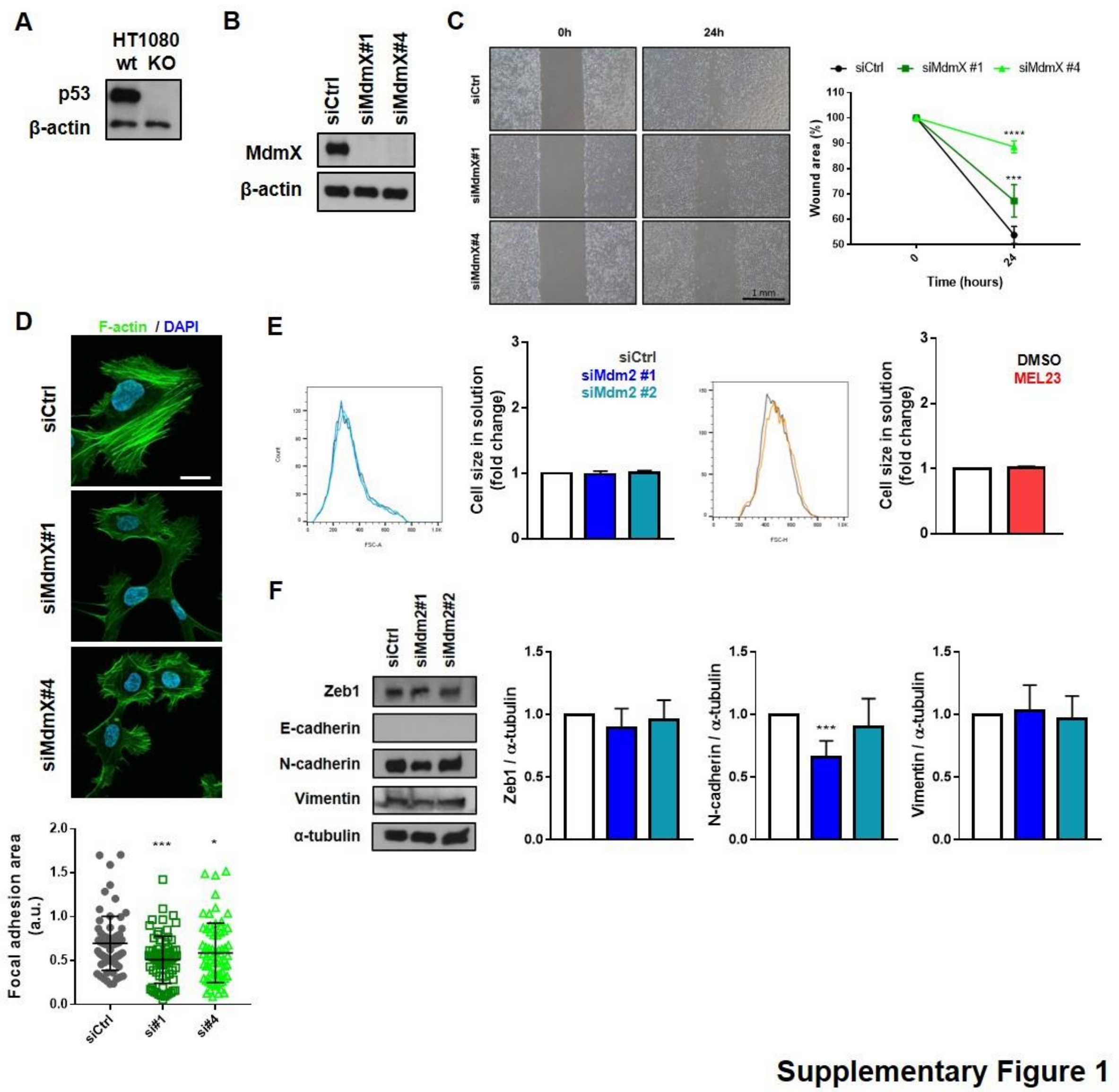

**Supplementary Fig. 1 Functional aspects of ablation of Mdm2 and MdmX in HT1080 p53KO cells. (A)** p53 protein levels in HT1080 parental (p53 wild-type, wt) and HT1080 p53KO cells. β-actin was used as loading control. **(B-D)** HT1080 p53KO cells were silenced for MdmX using siRNAs. (B) Protein levels of MdmX after transfection. β-actin was used as loading control. (C) Cell migration assay. Quantification and representative images of wound scratch migration assay. Scale bar equivalent to 1 mm. (D) Representative images of morphology of cells attached to collagen coated coverslips and quantification of cell area after silencing of MdmX. Scale bar equivalent to 50 µm. **(E)** Representative histogram and quantification of

HT1080 p53KO cell size in solution in response to Mdm2 knockdown or MEL23 treatment (7 µM) for 24 h. **(F)** Protein levels of EMT markers in HT1080 p53KO cells treated with control (siCrtl) or Mdm2 siRNAs. α-tubulin was used as loading control.

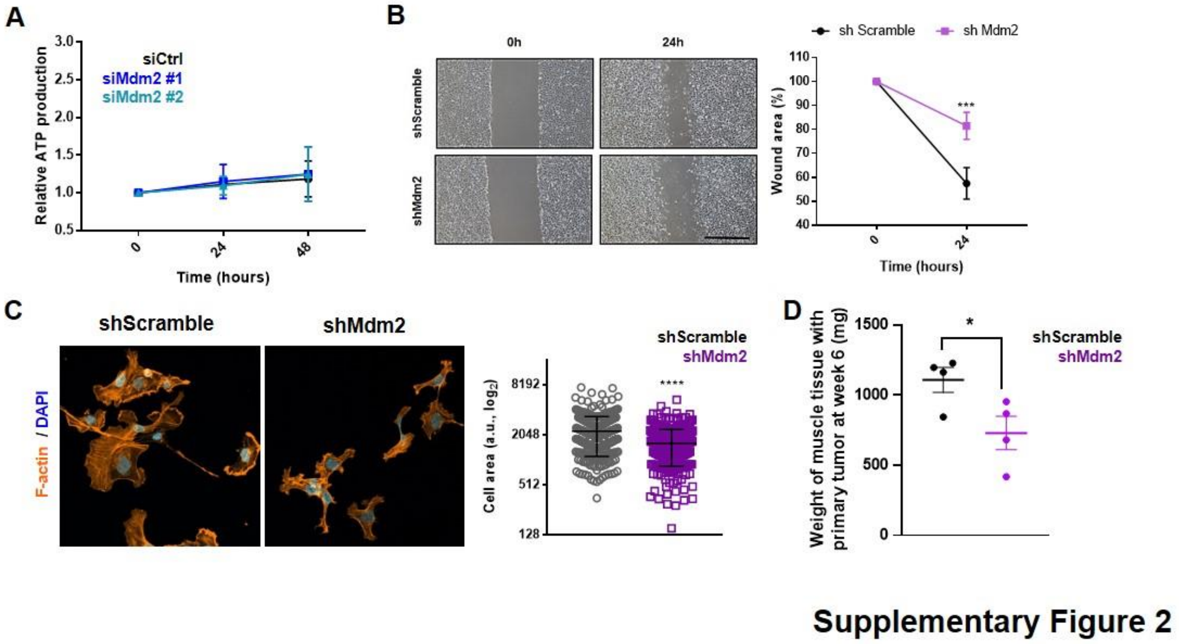

**Supplementary Fig. 2 Mdm2 depletion in growth and metastasis-related processes. (A)** Effects of transient Mdm2 silencing on growth of tumor spheroids of HT1080 p53KO cells measured by CellTiter Glo 3D. **(B)** Cell migration assay. Quantification and representative images of wound scratch migration assay using shScramble and shMdm2 stable GFP + cell lines. Scale bar equivalent to 1 mm. **(C)** Representative images (left) and quantification (right) of morphology of HT1080 p53KO shMdm2 and shScramble stable cell lines. Scale bar is equivalent to 50 µm. **(D)** Quantification of tumor weight at week 6 in the orthotopic mouse model. Experiments shown represent mean ±SD of at least 3 biological replicates, *p<0.05, **p<0.01, ****p<0.0001.

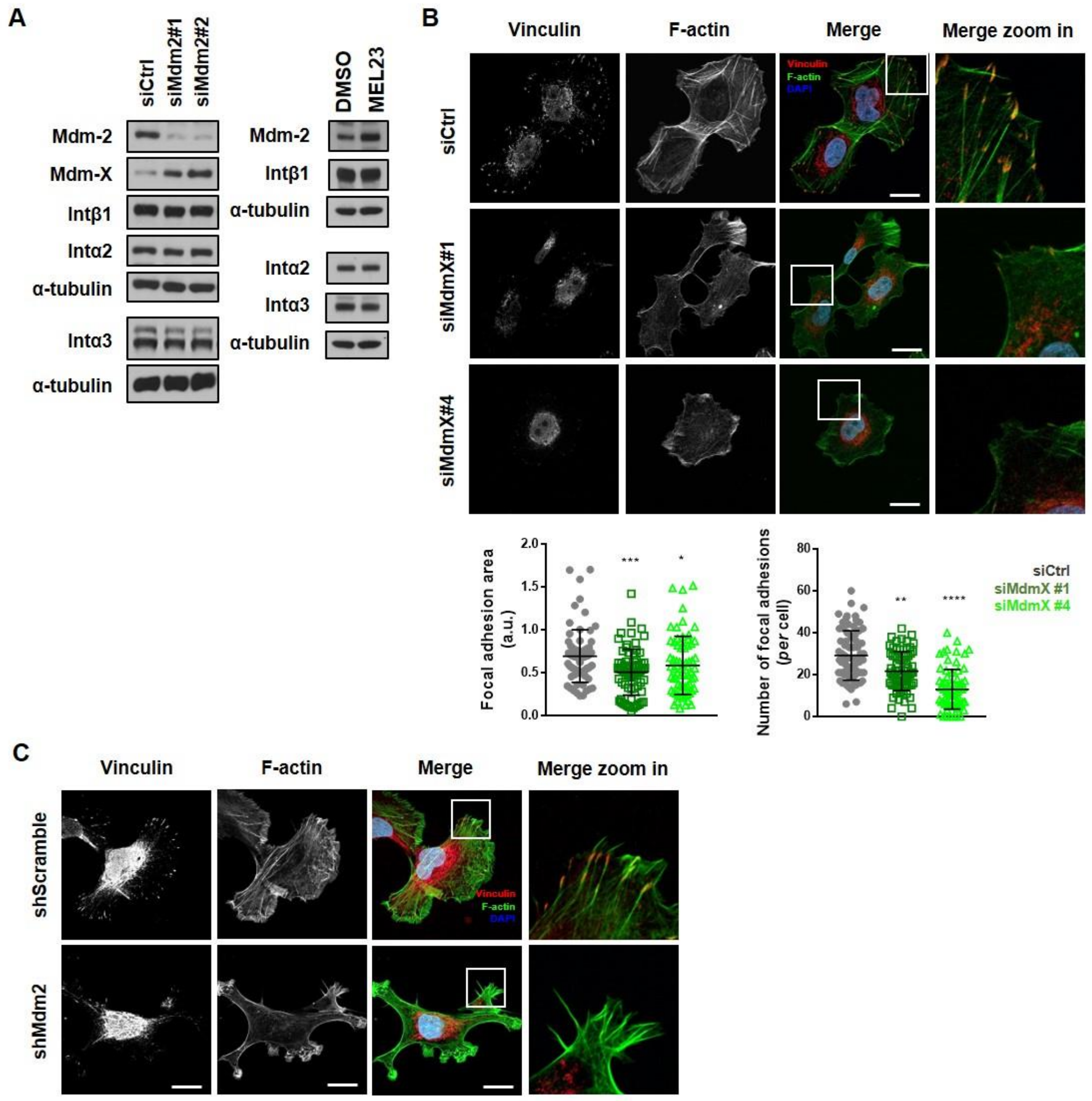

Supplementary Figure 3

**Supplementary Fig. 3 Changes in FA formation in response to Mdm2 loss. (A)** Protein levels of key expressed integrins in HT1080 p53KO cells in response to Mdm2 knockdown by siRNA (left) or MEL23 treatment (7 μM) for 24 h (right). α-tubulin was used as loading control. **(B-C)** Immunofluorescence showing FA foci by vinculin staining (red), stress fiber formation by phalloidin staining (green), and nuclei by DAPI staining of DNA (blue). (B) Representative images and quantification of FA parameters of HT1080 p53KO cells silenced for MdmX using siRNAs. (C) Representative images of HT1080 p53KO cells stably expressing shRNAs against Mdm2. Scale bar equivalent to 20 µm. Experiments shown represent mean ±SD of at least 3 biological replicates.

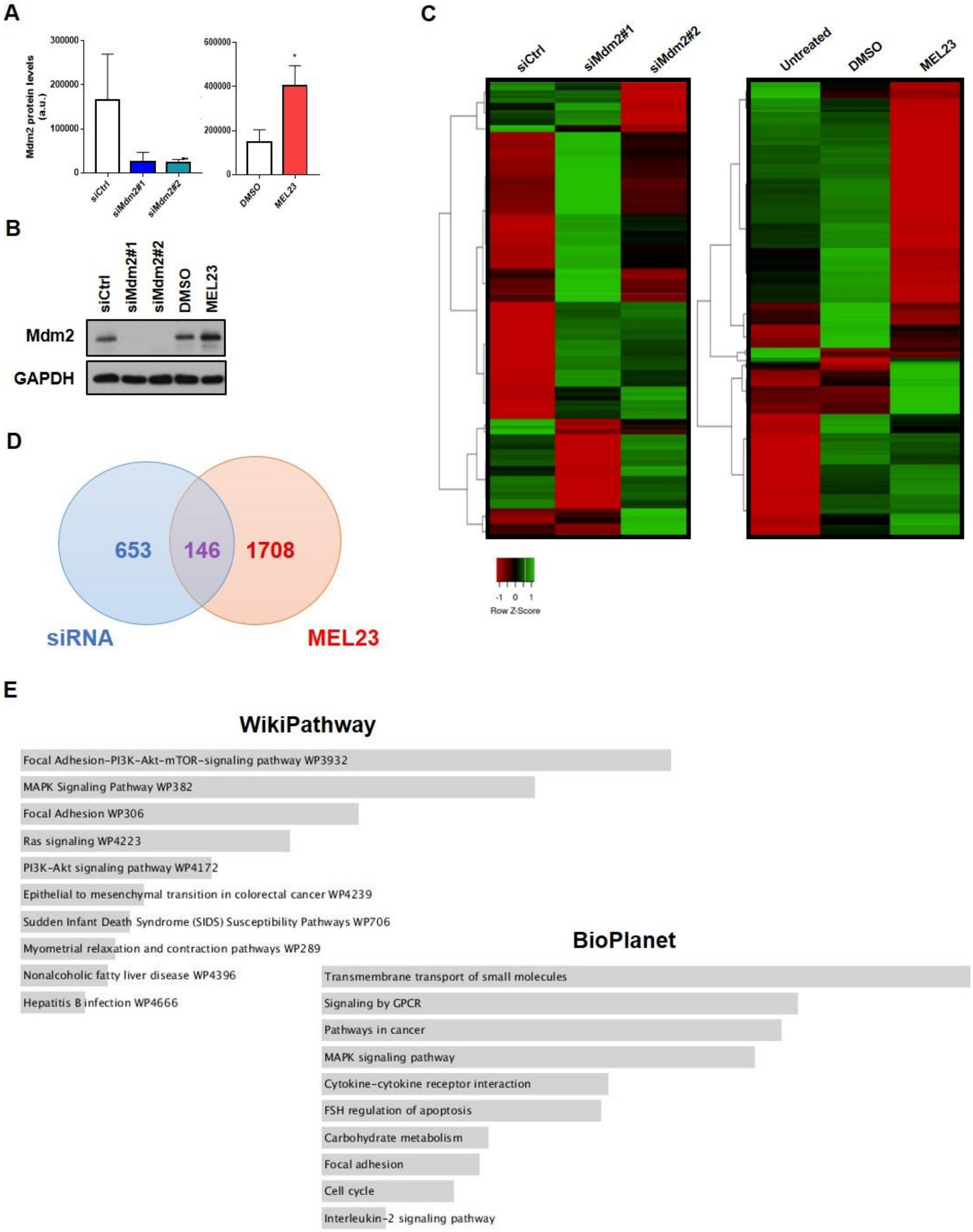

**Supplementary Figure 4**

**Supplementary Fig. 4 Proteomic analysis of HT1080 p53KO cells in response to Mdm2 inhibition using siRNAs and pharmacological inhibitor MEL23. (A-B)** Validation of conditions by quantification of Mdm2 expression in proteomics samples through (A) mass spectrometry and (B) immunoblot. GAPDH was used as loading control for the immunoblot. **(C)** Heatmaps of significantly differentially expressed proteins in the conditions indicated at top. **(D)** Venn diagram of proteins significantly differentially expressed only after transfection with siRNAs against Mdm2, or after treatment with 7 µM MEL23 or significantly changed by both strategies. **(E)** Pathway analysis of the 146 proteins significantly differentially expressed in both siRNA and drug groups analyzed in WikiPathway and BioPlanet databases. The proteomic analysis was performed with three independent biological replicates, *$p<0.05$.

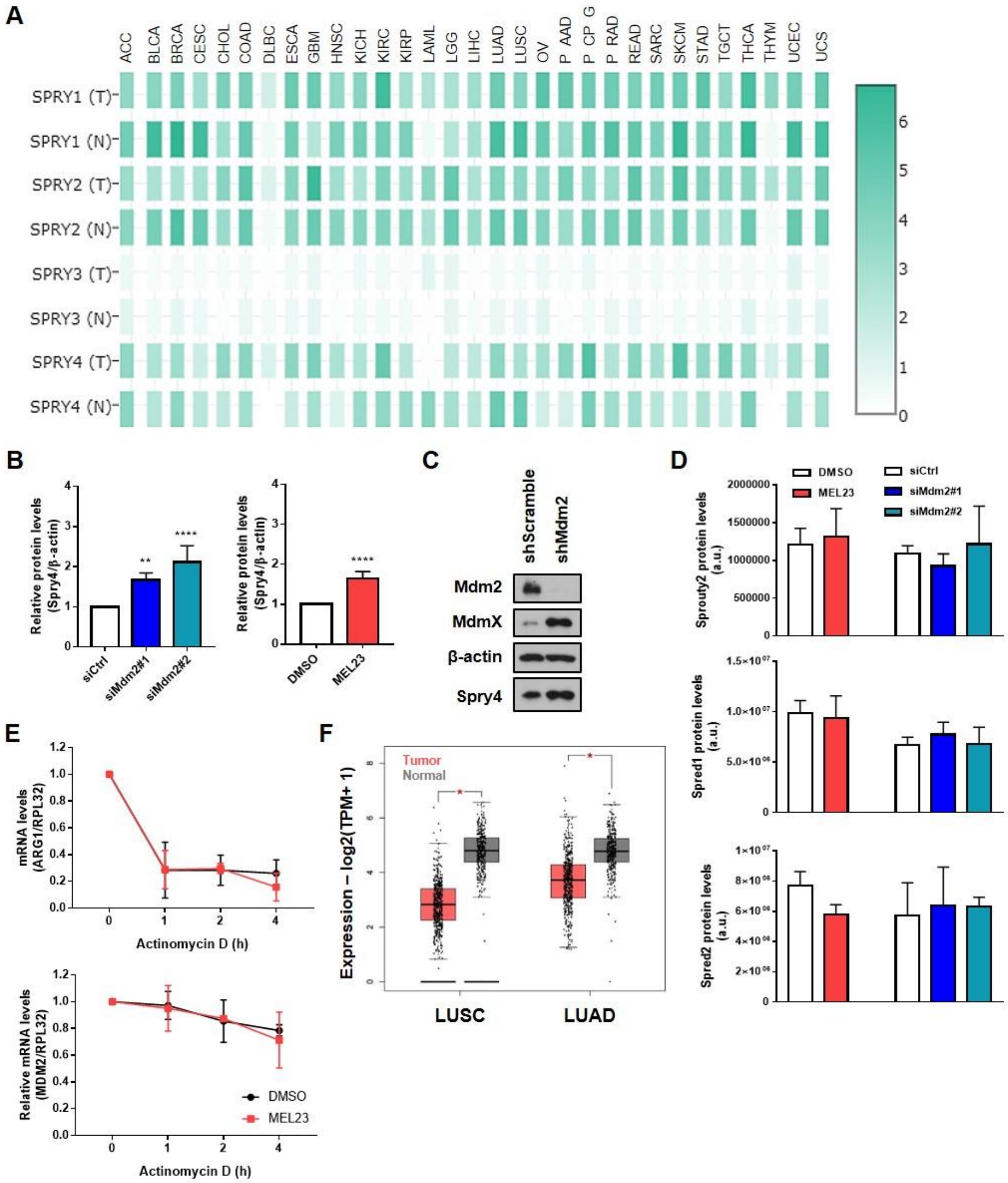

**Supplementary Figure 5**

**Supplementary Fig. 5 Sprouty expression in response to Mdm2 knockdown in different cell lines and patients' samples. (A)** Expression of Sprouty family members in patient samples from TCGA and GTEX databases. **(B)** Quantification of protein levels of Spry4 measured by densitometry of immunoblots in HT1080 p53KO cells silenced for Mdm2 (left) or treated with MEL23 (right). **(C)** Expression of Spry4 in response to stable Mdm2 knockdown in HT1080 p53KO cells. **(D)** Quantification of three Sprouty family members expressed in HT1080 cells in response to treatment with MEL23 or Mdm2 KD measured by MS proteomics analysis. **(E)** Half-life of ARG1 mRNA, as an example of a short-lived mRNA, and MDM2, as an example of a more stable mRNA, in cells treated with 7 µM MEL23 or vehicle in response to actD treatment. **(F)** TCGA and GTEX data analysis of Spry4 expression in normal lung and in lung carcinomas in patient samples. Experiments shown represent mean ±SD of at least 3 biological replicates, *p<0.05, **p<0.01, ****p<0.0001, n.s.: not significant.

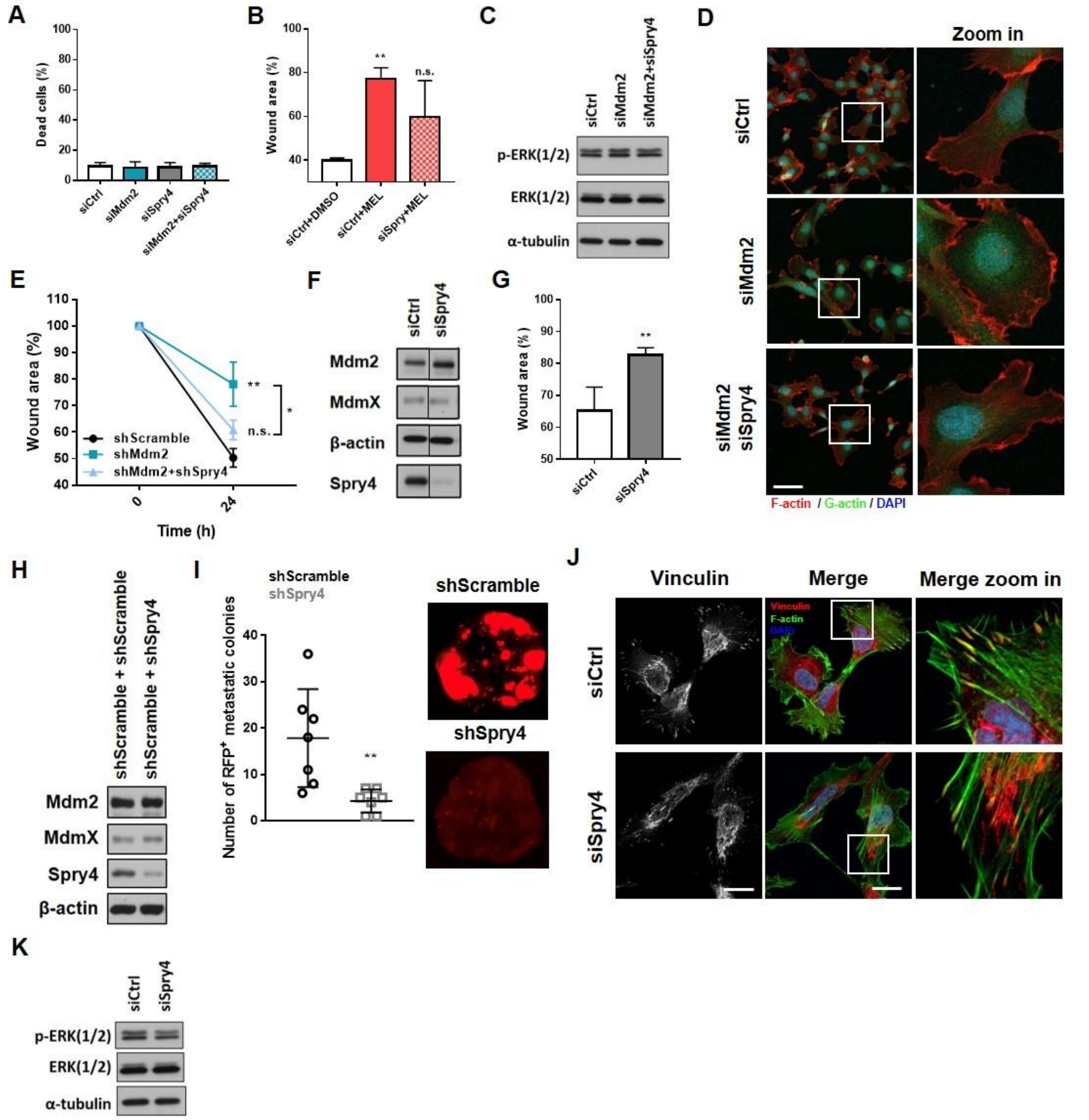

**Supplementary Figure 6**

**Supplementary Fig. 6 Experiments with Mdm2 and/or Spry4 siRNAs or with MEL23 treatment to test cell parameters related to Figure 5 and 6. (A)** Cell viability of HT1080 p53KO cells in response to Mdm2 knockdown as measured by Trypan Blue. **(B)** Quantification of migration of HT1080 p53KO cells silenced or not for Spry4 in the presence of absence of Mdm2 inhibitor MEL23. **(C)** Levels of ERK phosphorylation in response to knockdown of Mdm2 alone or co-knockdown of Mdm2 and Spry4 in HT1080 p53KO cells. **(D)** Actin polymerization staining. Representative images of fluorescent staining of F- (red) and G- (green) actin in HT1080 p53KO cells silenced for Mdm2 alone or with double knockdown of Mdm2 and Spry4. DAPI shows nuclei staining. **(E)** Cell migration assay. Quantification and representative images of wound scratch migration assay using shScramble, shMdm2 or shMdm2+shSpry4 stable cell lines. Scale bar equivalent to 1 mm. **(F-K)** HT1080 p53KO cells were silenced for Spry4 using a pool of si- or shRNAs. (F) Protein levels of Spry4 as well as Mdm2 and MdmX after transfection. α-tubulin was used as loading control. (G) Quantification of wound scratch migration assay of Spry4-depleted cells. (H) Protein levels of Spry4, Mdm2 and MdmX in HT1080 p53KO cells stably expressing shRNAs against Spry4. β-actin was used as loading control. (I) Analysis of metastatic burden *in vivo* using tail-vein model. Representative images and quantification of metastatic foci in the lungs using after 8 weeks of injection. (J) Immunofluorescence showing FA foci by vinculin staining, stress fiber formation by phalloidin staining, nuclei are detected by DAPI staining of cells transiently transfected with a pool of siRNAs against Spry4 or siCtrl. Scale bar equivalent to 20 µm. (K) Levels of ERK phosphorylation in response to transient knockdown of Spry4. Experiments shown represent mean ±SD of at least 3 biological replicates, *p>0.05, **p<0.01, n.s.: not significant.
